# Transcriptional Interference Gates Monogenic Odorant Receptor Expression in Ants

**DOI:** 10.1101/2025.08.21.671318

**Authors:** Giacomo L. Glotzer, P. Daniel H. Pastor, Daniel J. C. Kronauer

**Affiliations:** Laboratory of Social Evolution and Behavior, The Rockefeller University, New York, NY 10065, USA; Howard Hughes Medical Institute, New York, NY 10065, USA

## Abstract

Communication is crucial to social life, and in ants, it is mediated primarily through olfaction. Ants have more odorant receptor (OR) genes than any other group of insects, generated through tandem duplications that produce large genomic arrays of related genes. However, how olfactory sensory neurons produce a single functional OR from these arrays remains unclear. In ants, only mRNA from one OR in an array is exported into the cytoplasm, while upstream genes are silent and transcripts from downstream genes remain nuclear. Here, we show that non-canonical readthrough transcription in the downstream direction generates non-translated transcripts. We also find that OR promoters are bidirectional, producing antisense long non-coding RNAs that appear to suppress the expression of upstream genes. Finally, we present evidence that this regulatory architecture is conserved across ants and bees, suggesting that this mechanism for functionally monogenic OR expression is widespread in insects with expanded OR repertoires.

## INTRODUCTION

Almost all organisms rely on chemosensation to sample and interpret cues from the environment. The ability to perceive different compounds is dictated by a species’ chemoreceptor repertoire. Odorant receptors (ORs), which mediate the sense of smell, are a major class of chemoreceptors. While vertebrate ORs are GPCRs that signal via secondary messengers, insect ORs function as ligand-gated cation channels and form heterotetramers of a conserved coreceptor, Orco, and one of many tuning ORs.^1^

Eusocial insects, like ants, some wasps, some bees, and termites, rely heavily on chemosensation, as they use pheromone communication to coordinate their sophisticated social and collective behaviors. In ants, this is reflected by expanded OR repertoires.^2–4^ While the fruit fly *Drosophila melanogaster* has roughly 60 ORs, ants possess up to 687 ORs.^5^ When Orco function is disrupted, ants become essentially asocial, illustrating the importance of ORs in mediating ant sociality.^6,7^

A longstanding question in sensory biology is how olfactory sensory neurons (OSNs) select which receptor from their repertoire to express.^8–14^ In fruit flies and mice, each OSN generally expresses a single OR, and the axons of OSNs expressing the same receptor converge on shared regions, known as glomeruli, in the primary olfactory centers of the brain.^15–19^ Fly OSNs rely on a combinatorial code of transcription factors to dictate OR expression and axon guidance,^20,21^ whereas mouse OSNs exhibit semi-random OR choice.^10,22,23^

Our understanding of how OSNs choose which OR to express in insects other than *Drosophila* remains limited. Importantly, many insects differ from *Drosophila* in the genomic organization of ORs, suggesting that they might use distinct mechanisms for receptor choice. While in *Drosophila* almost all ORs occur as isolated singletons,^24^ many ORs in other insects are arranged in tandem arrays, clusters of genes aligned on the same DNA strand that are hotspots for gene duplication.^25–34^ Ant tandem arrays can be particularly large and numerous. The clonal raider ant, *Ooceraea biroi*, for example, which has 503 ORs, has over 40 tandem arrays of 2-82 ORs distributed across 13 of its 14 chromosomes.^27^

Within insect tandem arrays, coexpression of multiple ORs is common.^35–37^ Specifically, ant OSNs express a primary “chosen” OR alongside additional ORs located downstream in the same tandem array.^35^ Importantly, only RNA from the chosen OR is exported into the cytoplasm where it can be translated into protein, while RNA from downstream ORs remains sequestered in the nucleus.^35^ Furthermore, single nucleus multiomic sequencing (RNA-seq and ATAC-seq) of honeybee (*Apis mellifera*) OSNs revealed that a single active promoter within a tandem array can drive the coexpression of multiple downstream ORs.^37^ However, beyond these initial insights, the gene regulatory mechanisms that dictate functionally monogenic receptor expression at insect OR tandem arrays remain unknown.

Here, we show that bidirectional transcription from a single promoter enhances selectivity in receptor expression. Non-canonical transcriptional readthrough ensures that downstream genes do not produce transcripts with protein-coding potential. ORs located upstream of the chosen OR are transcriptionally inhibited by an antisense long non-coding RNA (lncRNA) that originates from the same promoter as the chosen OR. This mechanism can explain transcriptional patterns at the edge of tandem arrays and in rare cases where ORs within a tandem array have been inverted. Collectively, we interpret this extensive transcriptional activity as a regulatory filter that ensures monogenic OR protein production in ants and other hymenopteran insects. We propose that this mechanism may facilitate the integration of newly duplicated genes into the olfactory system and therefore contribute to the rapid expansion of OR repertoires.

## RESULTS

### OR Expression Affects the Transcription of Downstream Non-OR Genes

In principle, the mechanism that drives the expression and nuclear sequestration of ORs downstream of the chosen OR could be a function of the local transcriptional landscape or encoded in the OR genes themselves. To test these possibilities, we examined the expression of non-OR genes that are either adjacent to, or nested within, OR tandem arrays. We first quantified the expression of genes directly flanking each tandem array using published snRNA-seq data.^35^ Surprisingly, many of these flanking genes showed enrichment in the cluster of cells that expressed ORs in the corresponding tandem array. For non-OR genes downstream of tandem arrays, this enrichment was limited to genes located on the same DNA strand as the array (Figure 1A), whereas genes on the opposite strand exhibited no enrichment (Figure 1B). Even non-OR genes downstream of and in line with singleton ORs were specifically upregulated in cells that had chosen these ORs (Figure S1A and S1B), indicating that this phenomenon is not limited to tandem arrays. This expression pattern suggests that the transcriptional activity that drives downstream OR expression extends to non-OR genes.

**Figure 1.**
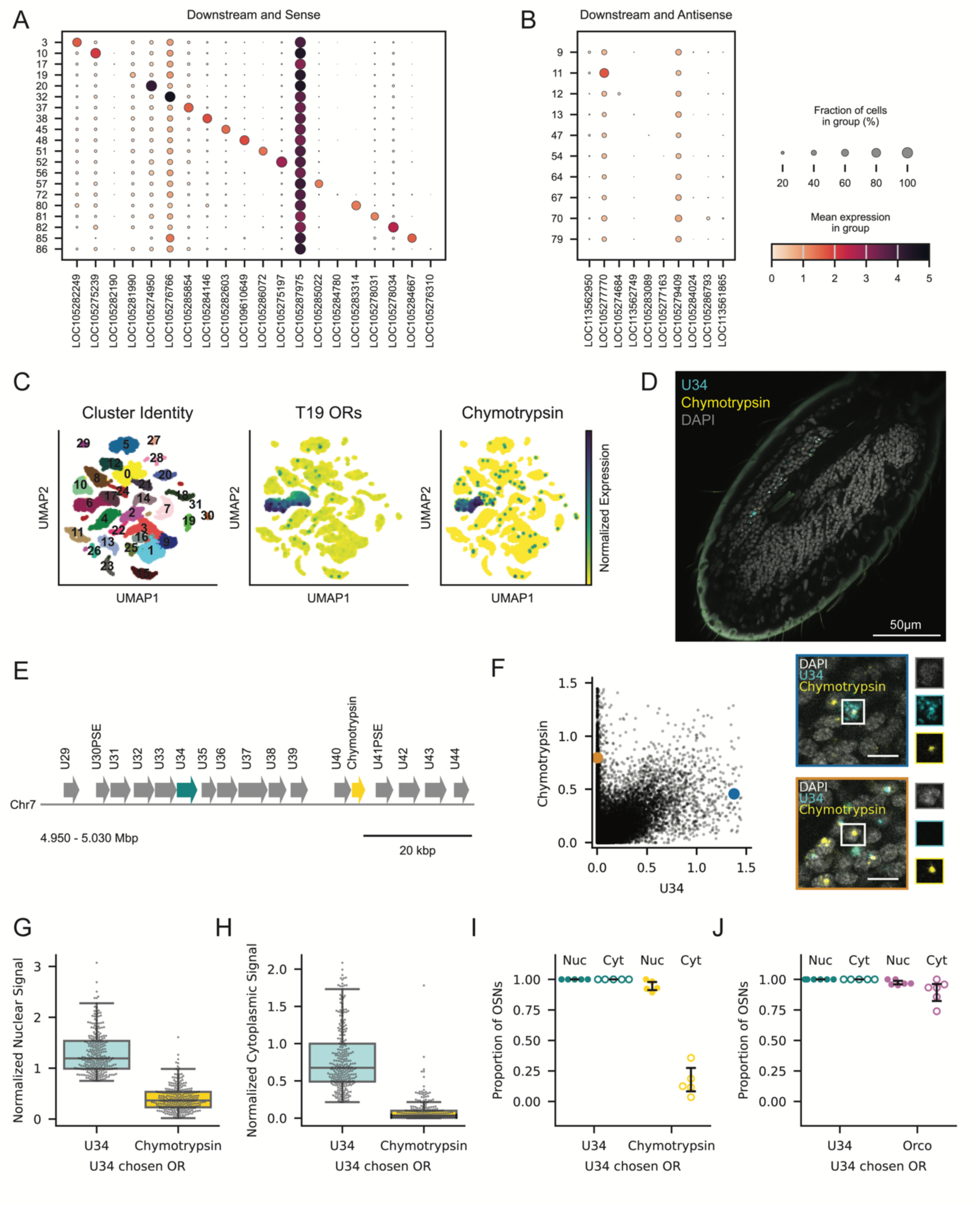
Coexpression of Genes Downstream of Chosen ORs Extends to Non-OR Genes. (A-B) Expression of non-OR genes downstream of OR tandem arrays on the same strand (A) or opposite strand (B) as the ORs. Column *n* corresponds to the downstream gene of the tandem array in row *n*. Cells are assigned a tandem array based on chosen OR expression, and a tandem array’s strand reflects the orientation of most ORs. Dot size corresponds to percentage of cells in each group that express a gene at a detectable level (>0) and dot color reflects the log-normalized expression level. (C) UMAP of antennal neurons colored by cluster (left), mean expression of T19 ORs (middle), and expression of chymotrypsin (right). (D) Slice from a confocal z-stack of an ant antennal club stained for U34 (cyan), chymotrypsin (yellow) and DAPI (grey). (E) Schematic of a subset of T19 highlighting the position of U34 (cyan) and the nested non- OR gene chymotrypsin (yellow). (F) Normalized nuclear signal for chymotrypsin vs. U34 in segmented OSN nuclei from n=5 antennae. Images with colored borders show the cells labeled with the corresponding colors, and each channel is shown individually to the right of each image. Blue: cell with cytoplasmic U34 and nuclear chymotrypsin. Orange: cell with cytoplasmic chymotrypsin only. Scale bars: 5 µm. (G-H) Normalized nuclear (G) and cytoplasmic (H) signal for U34 and chymotrypsin in 319 cells across n=5 antennae that express U34 as the chosen OR. Each boxplot shows the median and quartiles; the whiskers extend to 1.5 times the interquartile range. (I-J) Proportion of OSNs with U34 as the chosen OR per antenna (n=5) exhibiting U34 and chymotrypsin signal in the nucleus and cytoplasm. (J) Proportion of OSNs with U34 as the chosen OR per antenna (n=5) exhibiting U34 and Orco signal in the nucleus and cytoplasm. Error bars: 95% CI centered on the mean.

We also found that non-OR genes nested within tandem arrays undergo OR-mediated induction. Chymotrypsin (LOC105276652), a serine protease involved in digestion^38,39^ that has no known function in insect OSNs, is nested in tandem array T19 between OR U40 and the pseudogene U41. While chymotrypsin is not expressed in most OSNs, it is upregulated in the cluster of OSNs that express T19 ORs (Figure 1C). We conducted RNA-FISH staining for chymotrypsin and U34 (Figure 1D), an OR located 29 kbp upstream (Figure 1E). To quantify the RNA-FISH data for this and subsequent experiments, we trained a custom Cellpose 3.0 model^40^ to segment nuclei (Video S1) and cytoplasm (Video S2), and used consistent thresholds across all experiments to classify cells with nuclear and cytoplasmic transcripts (Methods). Across all OSNs, we found that cells expressing U34 generally also expressed chymotrypsin, whereas cells expressing chymotrypsin did not necessarily express U34 (Figure 1F). Because transcription can initiate anywhere along the array, we suspect that these chymotrypsin+/U34-cells had chosen an OR between U34 and U40. Cells that had chosen U34 had strong nuclear U34 and moderate nuclear chymotrypsin signal (Figure 1G). In contrast, the cytoplasmic intensity in these cells was high for U34 and very low for chymotrypsin, consistent with chymotrypsin RNA being sequestered in the nucleus (Figure 1H). We confirmed that 95% of these cells had chymotrypsin transcripts in the nucleus, but only 16% had transcripts in the cytoplasm, a level consistent with nonspecific background fluorescence (Figure 1I). Thus, while T19-expressing OSNs produce chymotrypsin transcripts, these transcripts likely do not yield functional protein. As a positive control, we confirmed that our signal segmentation pipeline detects cytoplasmic Orco transcripts in cells that express U34 (Figure 1J).

Finally, we examined the expression of two non-OR genes that border tandem arrays, identified in our analysis of flanking genes (Figure 1A). LOC105282603 and LOC105286072 are both uncharacaterized non-OR genes located directly downstream of T45 and T51, respectively (Figure S1C and S1D). Both genes are highly enriched in the cluster of cells expressing from their respective neighboring tandem array (Figure S1E). We found LOC105282603 coexpressed in 87% of cells that had chosen an upstream OR, with cytoplasmic localization in 6% of cases (Figure S1F). LOC105286072 was coexpressed in 100% of cells that had chosen an upstream OR, 52% of which also had cytoplasmic transcripts (Figure S1G and S1H). This demonstrates that non-OR genes located downstream and on the same strand as a tandem array are coexpressed in cells expressing an OR in that array, and transcripts from these non-OR genes can also undergo nuclear sequestration. The mechanism that drives downstream expression at ant OR tandem arrays thus extends beyond the boundaries of tandem arrays and operates independently of the protein products encoded by these genes.

At least two alternative models could explain these results. First, RNA polymerase II (RNAPII) might produce a continuous polycistronic transcript spanning the chosen OR and downstream genes. However, previous work using long-read mRNA sequencing and RNA-FISH suggests that each gene produces individual transcripts.^35^ Furthermore, mRNA from downstream genes is abundantly represented in 10X 3’ snRNA-seq data, which relies on oligo(dT) primers, implying that RNA from each downstream gene is also polyadenylated. An alternative possibility is that RNAPII terminates normally, but proximal genes produce transcripts due to leaky regulation. This scenario still does not explain why transcripts from downstream genes remain in the nucleus. The most parsimonious explanation involves a novel form of transcriptional readthrough, where a single RNAPII produces transcripts from several tandemly arrayed genes. Transcripts from each gene are cleaved and polyadenylated, but we suspect only transcripts from the first gene benefit from the addition of a 5’ cap, a key signal for nuclear export that is added immediately after transcription initiation.^41–43^

### Downstream Genes are Expressed as a Product of Transcriptional Readthrough

To test this hypothesis, we investigated whether RNAPII fails to terminate when transcribing OR genes, in which case we should be able to detect RNA from intergenic regions. We used an existing dataset of rRNA-depleted RNA-sequencing from whole *O. biroi* pupae,^44^ which includes nascent and non-polyadenylated transcripts. To compare the sequencing coverage of intergenic regions in tandem arrays with that of a comparable sample of paired non-OR genes, we sampled pairs of genes in the genome that were located on the same DNA strand with intergenic distances similar to those of typical OR gene pairs (Figure S2A; Methods). The mean coverage of OR exons was lower than that of non-ORs (Figure S2B), which is expected because ORs are expressed in a small fraction of cells in the whole pupa. We therefore normalized to the mean coverage of the upstream gene’s exons. Intriguingly, we found that OR intergenic regions had significantly higher relative coverage than intergenic regions between non-ORs (Figure 2A), consistent with the hypothesis that RNAPII fails to terminate transcription upon reaching the polyadenylation signal sequence (PAS) of OR genes.

**Figure 2.**
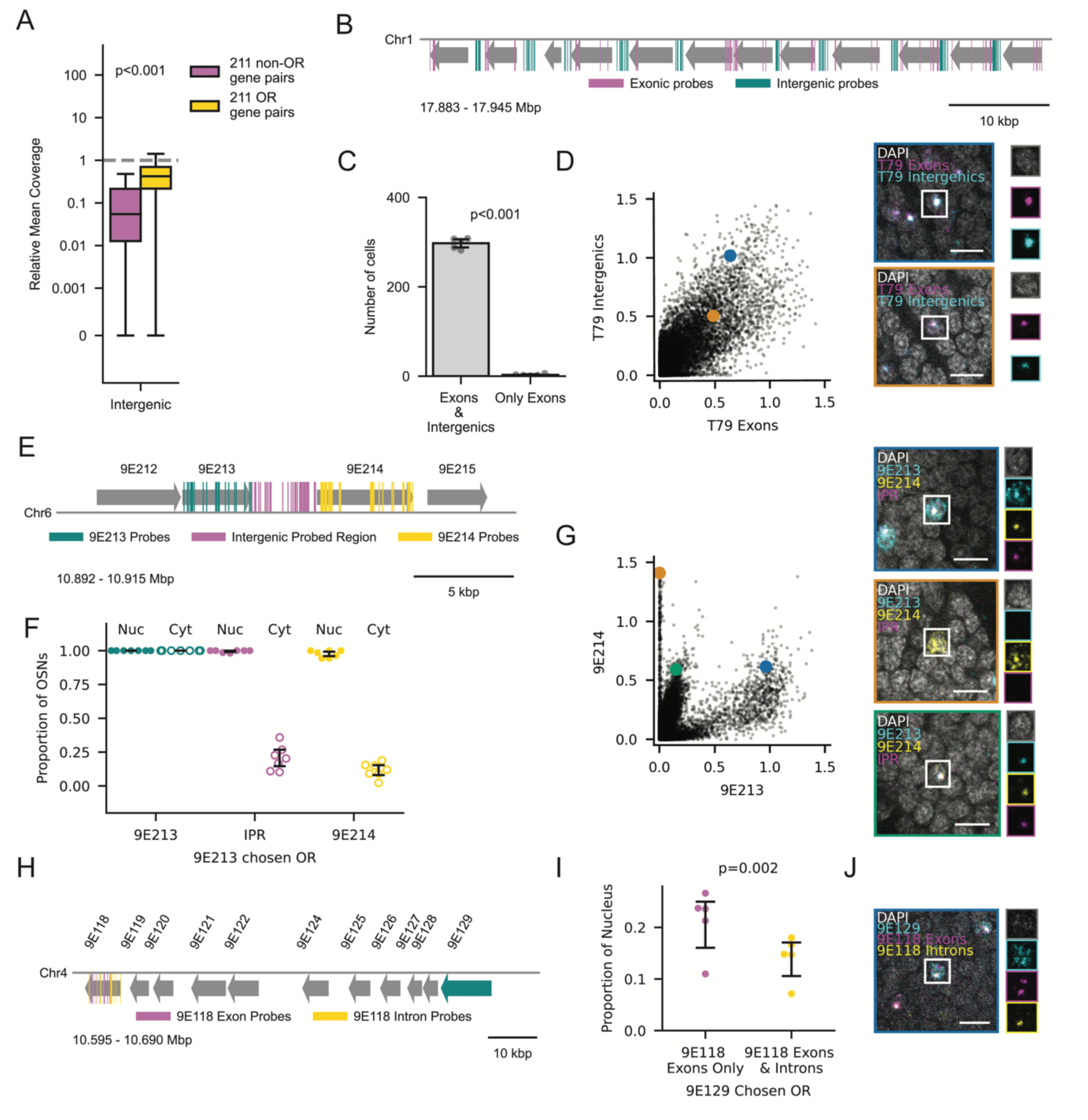
Intergenic Regions are Transcribed Along with Downstream Genes. (A) Relative coverage of intergenic regions for 211 non-OR (magenta) and 211 OR (yellow) gene pairs, normalized to the mean coverage of upstream exons. Each boxplot shows the median and quartiles; the whiskers extend to 1.5 times the interquartile range. The y-axis has a linear region from 0-0.001 and logarithmic scaling (base 10) outside this range. P-value from Wilcoxon rank-sum test. Dotted line at y=1. (B) Schematic of T79 with exon probe-binding regions in magenta and intergenic probe-binding regions in cyan. (C) Number of cells per antenna (n=5) with exonic and intergenic signal versus exon-only signal. P-value from two-sided t-test. (D) Normalized nuclear signal for T79 intergenics vs. T79 exons in segmented OSN nuclei from n=5 antennae. Images with colored borders show the cells labeled with the corresponding colors, and each channel is shown individually to the right of each image. Both images show example cells with both exonic and intergenic signal. (E) Schematic of a subset of tandem array T37 with probe-binding regions that target 9E213 (cyan), 9E214 (yellow) and the intergenic probed region (magenta). (F) Proportion of OSNs with 9E213 as the chosen OR per antenna (n=7) exhibiting 9E213, intergenic PR and 9E214 signal in the nucleus and cytoplasm. (G) Normalized nuclear signal for 9E214 vs. 9E213 in segmented OSN nuclei from n=7 antennae. Blue: cell with cytoplasmic 9E213, nuclear 9E214 and nuclear IPR. Orange: cell with cytoplasmic 9E214 only. Green: cell with nuclear 9E213, 9E214 and IPR. (H) Schematic of a subset of tandem array T45 with probe-binding regions that target 9E118 exons (magenta), 9E118 introns (yellow) and 9E129 exons (cyan). (I) Proportion of nucleus occupied by 9E118 exon signal alone and the overlap of exon and intron signal in cells with 9E129 as the chosen OR. (J) Example cell with cytoplasmic 9E129 and nuclear 9E118. Only some of the 9E118 exon signal overlaps with the 9E118 intron signal. Error bars: 95% CI centered on the mean. Scale bars: 5µm.

To validate that intergenic regions are transcribed with their respective coding regions, we designed FISH probes against T79 that target either the exons of all its genes or its intergenic regions (Figure 2B). T79 exhibits the characteristic staircase-like pattern of OR expression, where ORs downstream of the chosen OR are coexpressed (Figure S2C). The choice of T79 as an example array was based on long-read RNA sequencing data that suggested well-defined intergenic regions (Figure S2D). Antennae (n=5) contained 300 ±10 (range: 282-308) T79-expressing cells on average, with 99% of these cells labelled by both sets of probes (Figure 2C). In segmented nuclei, the T79 exon signal highly correlated with the intergenic signal (R=0.81, p<0.001) and the exon signal colocalized with the brightest intergenic signal (Figure 2D), suggesting that they label the same set of molecules.

Because downstream genes are also polyadenylated, we hypothesized that cleavage occurs co-transcriptionally but is insufficient to terminate transcription. We stained 9E213, 9E214 and the 3.3 kbp intergenic region (Figure 2E). In cells expressing 9E213 as the chosen OR, both the intergenic region and 9E214 were highly coexpressed, and their transcripts overlapped spatially and remained in the nucleus (Figure 2F and 2G). This indicates that cleavage occurs immediately after the 3’ end of 9E213 and suggests that the intergenic region and 9E214 are part of the same mRNA molecule. The 9E214 and intergenic transcripts colocalized strictly to a single region of the nucleus, where 9E213 signal was maximal, presumably the site of active transcription (Figure 2G). As expected, cells that expressed 9E214 as the chosen OR did not express 9E213 (Figure 2G and S2E).

For genes containing introns, splicing is a critical prerequisite for the nuclear export of transcripts.^45–47^ Therefore, we sought to determine whether transcripts from downstream ORs undergo proper splicing. We separately stained for 9E118 exons and introns, as well as 9E129, an OR located 65 kbp upstream (Figure 2H). We segmented the RNA-FISH signal as before and quantified the fraction of the nucleus occupied by 9E118 exon probes, which label nascent and mature transcripts, versus the region labeled by both 9E118 exon and intron probes, which label unspliced transcripts. In cells with 9E129 as the chosen OR, the exons occupied a larger area than the exon-intron overlap, suggesting that some degree of splicing does occur (Figure 2I). We confirmed visually that exonic and intronic signals do not perfectly overlap (Figure 2J). This indicates that transcripts from downstream ORs are spliced, and that another feature must be responsible for confining them to the nucleus.

Collectively, our results support the hypothesis that downstream OR expression is a product of a novel form of transcriptional readthrough in which transcripts from each gene are cleaved to produce monocistronic, spliced mRNAs. We surmise that, as in *A. mellifera,* a single *O. biroi* promoter can drive the expression of multiple downstream ORs.^37^ However, we argue against a polycistronic mode of transcription, and given that RNA-FISH of coexpressed ORs in the honeybee produces non-overlapping signal, we suspect the mechanism we describe here also occurs in bees. While the function of this non-canonical readthrough remains unknown, we suspect that it serves as a form of transcriptional interference, inhibiting the production of capped transcripts from downstream genes. Although this mechanism may explain why downstream ORs do not generate protein, an additional process must account for the silencing of ORs located upstream of the active promoter.

### Ant OSNs Produce Antisense RNA from OR Tandem Arrays

Returning to our previous analysis of flanking non-OR genes, we next looked at the genes located directly upstream of each tandem array. Intriguingly, we found the exact opposite pattern compared to downstream genes: Genes that were upstream and on the same strand exhibited minimal enrichment (Figure 3A), while genes upstream and on the opposite strand were highly upregulated in their respective clusters (Figure 3B). The same enrichment of antisense non-OR genes was present upstream of singleton ORs (Figure S3A and S3B). This pattern raises the question of whether these upstream antisense non-OR genes produce protein in this context.

**Figure 3.**
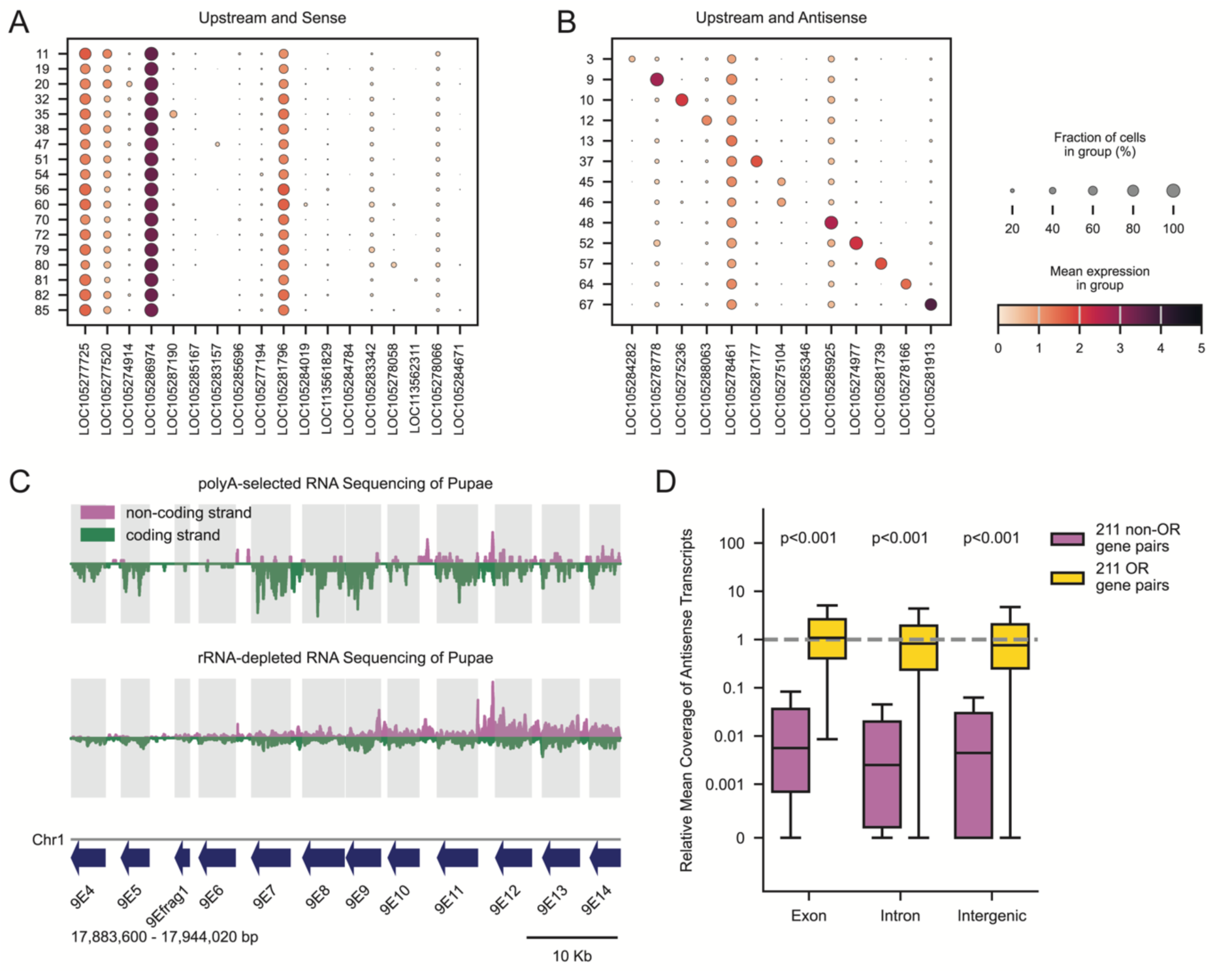
Tandem Arrays are Hotspots of Antisense lncRNAs. (A-B) Expression of non-OR genes upstream of OR tandem arrays on the same strand (A) or opposite strand (B) as the ORs. Column *n* corresponds to the upstream gene of the tandem array in row *n*. Cells are assigned a tandem array based on chosen OR expression, and a tandem array’s strand reflects the orientation of most ORs. Dot size corresponds to percentage of cells in each group that express a gene at a detectable level (>0) and dot color reflects the log-normalized expression level. (C) Strand-specific mRNA sequencing coverage (reads per nucleotide) across T79, using polyA-enriched RNA-seq (top) and rRNA-depleted RNA-seq (bottom) from whole pupae. Coding strand coverage (green) is mirrored across the x-axis for visualization. Gray boxes indicate annotated OR gene boundaries. (D) Relative antisense coverage of exons, introns, and intergenic regions for 211 non-OR (magenta) and 211 OR (yellow) gene pairs, normalized to the mean coverage of upstream exons on the sense strand. Each boxplot shows the median and quartiles; the whiskers extend to 1.5 times the interquartile range. The y-axis has a linear region from 0-0.001 and logarithmic scaling (base 10) outside this range. P-values from Wilcoxon rank-sum tests with Benjamini–Hochberg FDR correction. Dotted line at y=1.

We therefore investigated the expression of one of these genes using RNA-FISH. Krüppel homolog 1 (Kr-h1; LOC105275104), a transcription factor and effector of juvenile hormone signaling,^48^ is directly upstream of and antisense to T45 (Figure S4A), and its expression is specifically enriched in T45-expressing cells (Figure S4B). Kr-h1 was coexpressed in 74% of cells that had chosen a T45 OR (Figure S4C). Surprisingly, only 5% of these cells exhibited Kr-h1 transcripts in the cytoplasm (Figure S4C and S4D), a pattern associated with downstream genes.

Antisense transcripts, a class of lncRNA transcribed from the opposite DNA strand of a protein-coding gene, have emerged as powerful regulators of gene expression in eukaryotes.^49–54^ Two recent studies have highlighted the abundance of lncRNAs in insect OSNs,^55,56^ making them promising candidates for OR regulation.^57^ While annotated lncRNAs occur throughout the *O. biroi* genome, we noticed that OR loci are heavily enriched for antisense lncRNAs, with 27% of OR genes overlapping with annotated antisense lncRNAs, compared to only 6% of non-OR genes (Figure S4E). Accordingly, the number of antisense lncRNAs nested within tandem arrays scales with the number of ORs in each tandem array (Figure S4F). Because lncRNA annotations are often incomplete, we wondered whether lncRNAs could in fact be even more ubiquitous in OR tandem arrays.

To enhance our coverage of lncRNAs, which may lack a polyA tail,^58^ we revisited our dataset of rRNA-depleted RNA-seq and compared its coverage with that of a control dataset that relied on oligo(dT) primers.^44^ Compared to polyA-selected data, the rRNA-depleted dataset revealed a pronounced enrichment of reads mapping to the non-coding strand of OR tandem arrays (Figure S4G). An example tandem array illustrating this phenomenon is shown in Figure 3C.

To investigate whether the presence of antisense transcripts in the rRNA-depleted data was specific to OR genes, we returned to the same pairs of ORs and non-ORs and quantified the relative coverage of reads that align to the non-coding strand. Antisense transcripts are typically expressed at much lower abundance than sense transcripts,^59^ and indeed we found that non-OR gene pairs exhibited a ∼100-fold reduction in opposite-strand read coverage (Figure 3D). However, sense and antisense transcripts were approximately equally abundant at OR loci across exons, introns, and intergenic regions (Figure 3D). This indicates that non-coding antisense transcripts are pervasive across OR tandem arrays, and that many of these transcripts are missing in current genome annotations, possibly due to a lack of polyadenylation. In fact, all regions antisense to OR genes that currently lack an antisense lncRNA annotation have similar relative coverage to existing annotated lncRNAs nested in tandem arrays (Figure S4H), suggesting that polyadenylation is not necessary for the production of these lncRNAs.

### Antisense lncRNAs Emerge from Bidirectional OR Promoters

The high abundance of annotated and unannotated antisense transcripts at OR tandem arrays provokes the question of how their expression is regulated. To better understand the initiation sites of sense and antisense transcription, we analyzed capped small-RNA sequencing (csRNA-seq) data from whole adult ants. csRNA-seq captures transcriptional start sites (TSSs) at single nucleotide resolution and can better detect transient or unstable RNA than conventional RNA-seq.^60,61^ We found twin sense and antisense peaks upstream of dozens of ORs, including those with (Figure 4A) and without (Figure 4B) annotated lncRNAs. Importantly, the antisense peaks tend to be farther from the OR than the sense ones, preventing RNAPII collision (Figure 4C). These upstream peaks are prominent irrespective of whether an OR is in a tandem array or not (Figure 4D). Studies have shown that many promoters can exhibit bidirectional initiation even if elongation and maturation of transcripts is predominantly unidirectional.^62^ We therefore analyzed the same non-OR pairs from the rRNA-depleted RNA-seq analysis and found that, on average, 34% of these genes had bidirectional csRNA-seq reads (Figure 4E). Peaks were less common at ORs than non-ORs (Figure 4E), likely due to the low relative expression of ORs in whole adult tissue. This shows that bidirectional initiation is not limited to ORs, but that ORs stand out in their ability to promote antisense elongation and transcript production.

**Figure 4.**
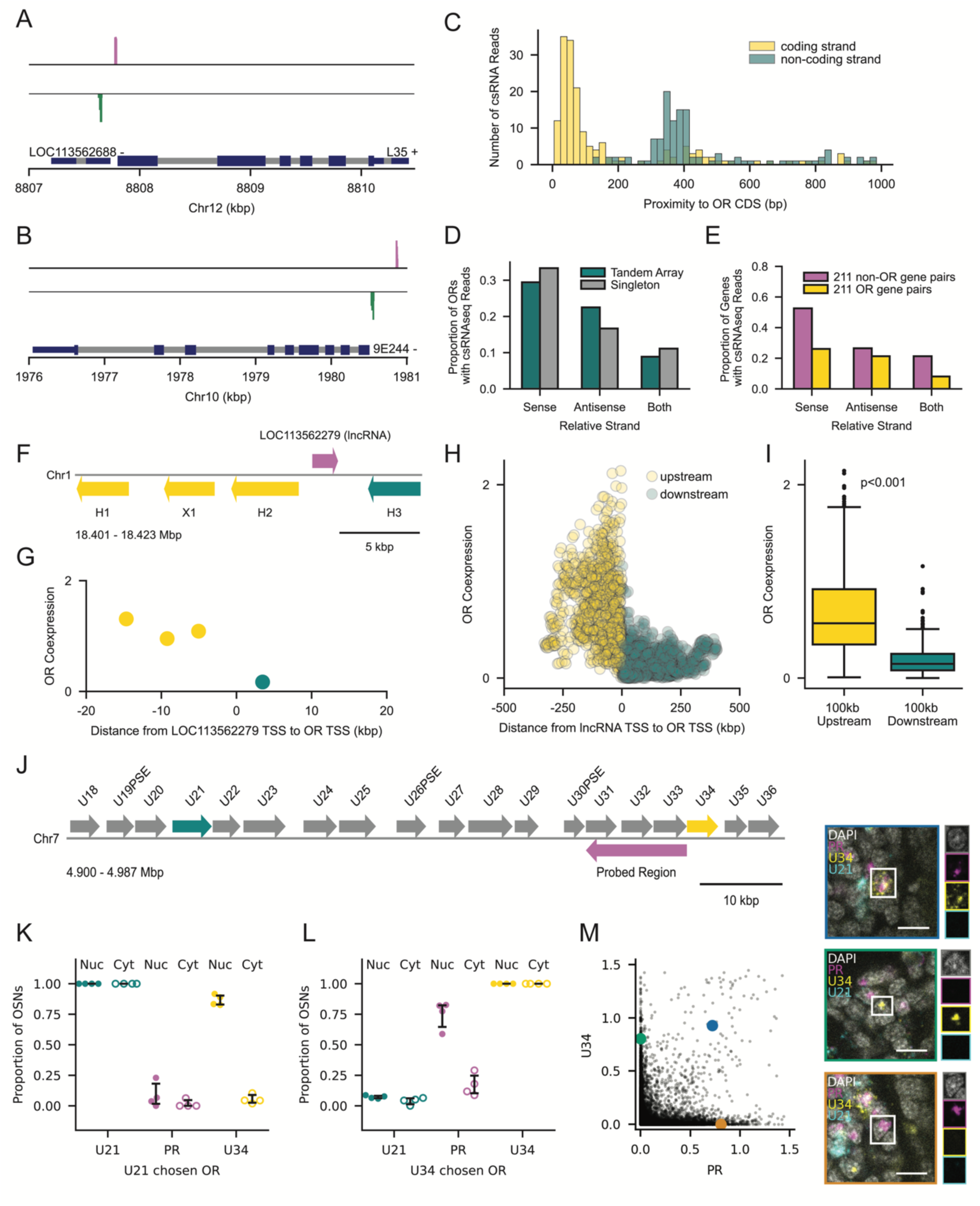
Antisense lncRNAs are Expressed from Bidirectional OR Promoters. (A-B) Bidirectional peaks from csRNA-seq upstream of L35 (A), an OR in T3 with an annotated lncRNA (LOC113562688), and 9E244 (B), an OR in T13. The strand orientation of each gene is indicated next to its name. Reads aligning to the sense and antisense directions are displayed in magenta and green, respectively. Gene models include exons (dark blue) and introns (grey), with coding sequences (CDS) in a thicker dark blue. (C) Histogram of the distance from each csRNA-seq read to the upstream OR CDS for reads on the same (yellow) and opposite (cyan) strand as the OR. Only reads within 1 kbp upstream of an OR’s first CDS are shown. (D) Proportion of ORs in tandem arrays (magenta) and singletons (grey) with csRNA-seq reads within 1 kbp upstream of the first CDS. The orientation of reads is given relative to the respective OR. (E) Proportion of 211 non-OR and 211 OR gene pairs with csRNA-seq reads within 1 kbp upstream of the TSS for non-ORs and first CDS for ORs. The sample is restricted to gene pairs arranged in tandem without antisense lncRNA annotations. (F) Schematic of T80, containing four ORs and a nested lncRNA (magenta), where the ORs are colored by their position relative to the lncRNA, either upstream (yellow) or downstream (cyan). (G) Mean log-normalized coexpression of each OR in T80 across nuclei with detectable expression (>0) of lncRNA LOC113562279 vs. TSS-TSS distance from LOC113562279. (H) Mean log-normalized coexpression of upstream (yellow) and downstream (cyan) ORs vs. the TSS-TSS distance from lncRNAs. Data include 70 unique lncRNAs embedded within tandem arrays and each dot represents a cell in which the corresponding lncRNA is detected. (I) Boxplot of log-normalized OR coexpression within 100 kbp upstream (yellow) or downstream (cyan) of antisense lncRNAs. Each boxplot shows the median and quartiles; the whiskers extend to 1.5 times the interquartile range. P-value from Wilcoxon rank-sum test. (J) Schematic of a subset of T19, highlighting ORs U21 (cyan), U34 (yellow), and the probed region (PR) targeting a putative antisense lncRNA (magenta). (K-L) Proportion of OSNs with U21 (K) or U34 (L) as the chosen OR per antenna (n=4) exhibiting U21, PR and U34 signal in the nucleus and cytoplasm. Error bars: 95% CI centered on the mean. (M) Normalized nuclear signal for U34 vs. PR in segmented OSN nuclei from n=4 antennae. Images with colored borders reflect the cells labeled with the corresponding colors and each channel is shown individually to the right of each image. Blue: cell with cytoplasmic U34 and nuclear PR. Green: only nuclear U34. Orange: only nuclear PR. Scale bars: 5 µm.

Although we suspect that antisense transcripts are ubiquitous within ant OR tandem arrays, our snRNA-seq analysis only includes reads that map to the 74 annotated lncRNAs nested within tandem arrays. For each cell expressing these lncRNAs, we plotted the mean expression level of nearby ORs against the genomic distance of these ORs from the lncRNA. An example is shown in Figures 4F and 4G, and aggregate data are shown in Figures 4H and 4I. This analysis revealed that ORs immediately upstream of lncRNAs are expressed at significantly higher levels than ORs immediately downstream of the lncRNA (Figure 4H and 4I), suggesting that these lncRNAs, along with transcripts from the upstream OR, originate from a single active promoter region that produces bidirectional transcriptional activity.

Because the vast majority of lncRNAs in the *O. biroi* genome currently appear to be unannotated, we aimed to verify their existence and to further investigate their coexpression with upstream ORs. First, we probed for a putative lncRNA in T19 by targeting a 12 kbp region antisense to U31-U33 (Figure 4J). In cells expressing the upstream U21 as the chosen OR, signal from the probed region was absent, whereas U34 RNA was present but restricted to the nucleus (Figure 4K). In contrast, cells with the downstream U34 as the chosen OR lacked U21 transcripts but coexpressed the probed region (Figure 4L). Transcripts from the probed region also remained in the nucleus (Figure 4L), a feature common to lncRNAs.^51^ Some cells also expressed U34 alone or the probed region alone (Figure 4M), which are likely cells with chosen ORs upstream and downstream of U34, respectively.

We then studied T70, a minimal tandem array consisting of only two ORs, Q1 and R2 (Figure S5A). Because of its high similarity to R3, a singleton on a different chromosome, we were unable to design probes unique to R2. Of the cells that had chosen Q1, 98% were labeled by R2/3 probes, indicating the presence of R2 transcripts (Figure S5B). In contrast, of the cells that had chosen R2/3, 71% coexpressed a putative antisense lncRNA antisense to Q1 (Figure S5C). The subset of cells that did not coexpress the putative lncRNA were likely R3-expressing cells (Figure S5D).

To ascertain whether this pattern of lncRNA expression is general to ant ORs, we looked for lncRNAs upstream of singleton ORs and confirmed coexpression where lncRNAs are annotated (Figure S4E and S4F) and unannotated (Figure S4G-I). Together, these results suggest that every OR gene in the ant genome, including singleton ORs, has a bidirectional promoter region that produces an antisense lncRNA in cells where the respective OR is expressed as the chosen OR.

Lastly, we investigated the length of these lncRNAs. For comparison, we knew that transcription in the sense direction can extend >100 kbp downstream (Figure S5J). Although the annotated lncRNAs can appear short (Figure 4F), we hypothesized that many mapped reads may represent transcription initiation much further upstream, as our quantification of coexpression suggests (Figure 4H). Using RNA-FISH, we found that 85% of cells produced a lncRNA >30 kbp in length (Figure S5K and S5L), and 61% produced a lncRNA >100 kbp in length (Figure S5M and S5N). Our results suggest that bidirectional transcription from a single OR promoter region can result in polymerase activity >100 kbp in both directions.

### Antisense lncRNA Expression Negatively Correlates with the Expression of Upstream ORs

While these experiments verify the existence of ubiquitous antisense lncRNAs at OR tandem arrays, their function remains unclear. We hypothesized that lncRNAs inhibit the transcription of ORs upstream of the active promoter, which may be prone to “off-target” transcription due to their spatial proximity. We revisited our snRNA-seq data and identified OSNs expressing a lncRNA upstream of their respective chosen OR. For each OSN, we correlated the expression level of the lncRNA with ORs flanking the chosen OR. For example, when we examined the cells with 9E300 as the chosen OR, we found that the expression of an upstream antisense lncRNA (LOC113562279) was negatively correlated with the expression of the upstream OR and positively correlated with the expression of the downstream OR (Figure 5A and 5B).

**Figure 5.**
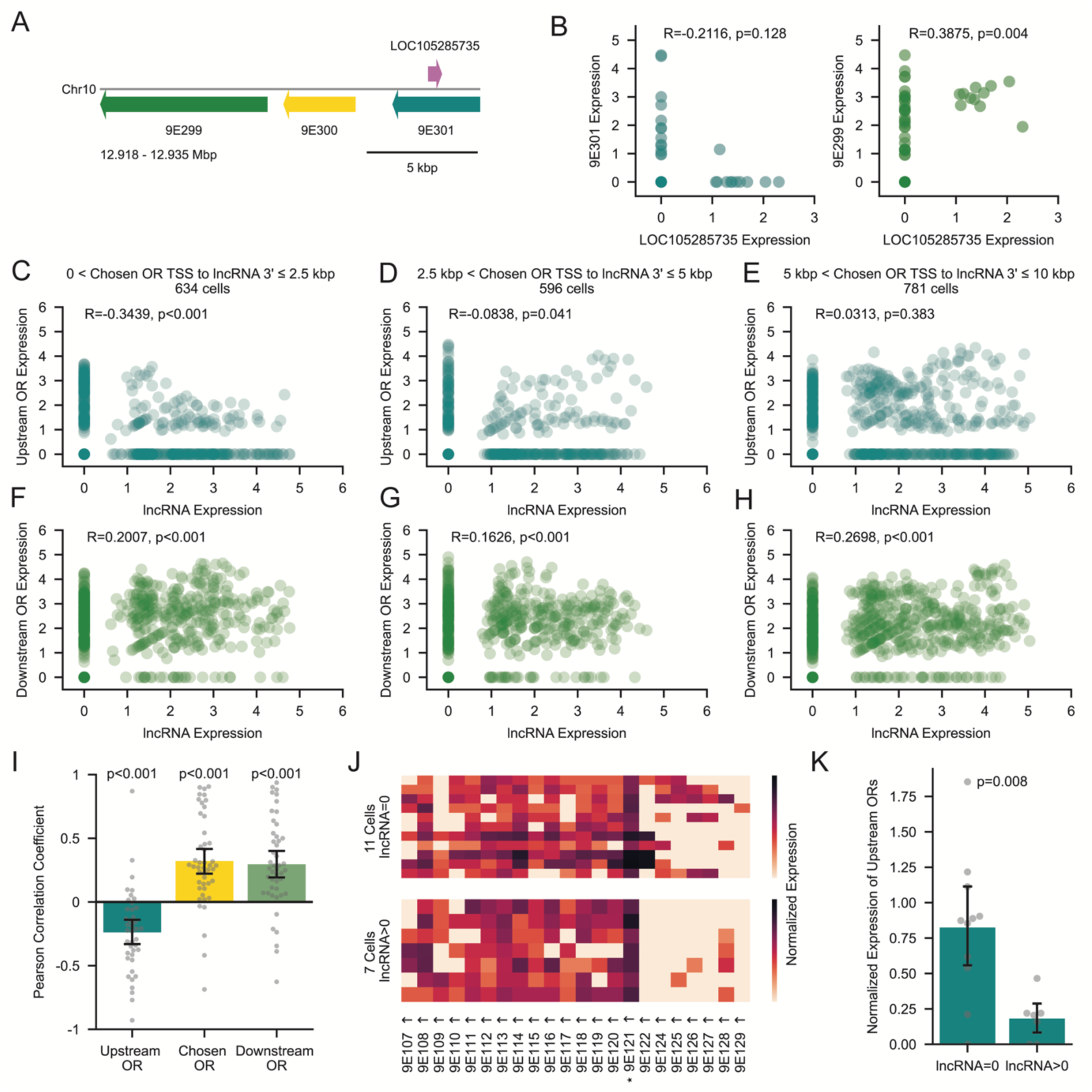
Antisense lncRNAs Prohibit Expression of Upstream ORs. (A) Schematic of a subset of T17 showing three ORs, 9E299 (green), 9E300 (yellow) and 9E301 (cyan) and the antisense lncRNA LOC113562279 (magenta). (B) Log-normalized expression of 9E301 versus LOC113562279 (left) and E299 vs. LOC113562279 (right), in cells where 9E300 is the chosen OR. (C-H) Scatterplots of log-normalized lncRNA versus upstream OR (C-E) or downstream OR (F-H) expression, split by the genomic distance from the chosen OR TSS to the 3’ end of the nearest upstream lncRNA: 0-2.5 kbp (C, F), 2.5-5 kbp (D, G), and 5-10 kbp (E, H). Pearson correlation coefficients and p-values are indicated above each plot (B-H). (I) Mean correlation of each lncRNA and either upstream ORs (cyan), chosen ORs (yellow), or downstream ORs (green). The 3’ end of each lncRNA is within ≤5 kb of the chosen OR TSS. P-values from one-sample t-tests against zero. (J) Heatmaps of log-normalized expression of all ORs in T45 for cells with 9E121 (*) as the chosen OR. Split by absent (top) or detectable (bottom) expression of the antisense lncRNA LOC109611203 located 2 kbp upstream of 9E121. Arrows indicate strand orientation. (K) Mean expression of upstream genes 9E122-129 for all cells in (J) split by absent or detectable expression of the antisense lncRNA LOC109611203. P-value from Wilcoxon rank-sum test. Error bars: 95% CI centered on the mean.

Aggregating these data across cells that express a chosen OR ≤2.5 kbp from a lncRNA, we confirmed that the expression levels of the lncRNA and upstream OR are negatively correlated (Figure 5C). At these small distances, OSNs exhibit a switch-like behavior, with 85% of cells showing either upstream OR expression and no lncRNA expression, or vice versa (Figure 5C). This correlation weakens in cells where the window is shifted to >2.5 kbp and ≤5 kbp (Figure 5D) and subsides entirely once the window increases to >5 kbp and ≤10 kbp (Figure 5E). For each of these distances, the correlation of lncRNA expression with downstream OR expression was positive, confirming that the activity of the bidirectional promoter region has opposite effects on upstream vs. downstream gene expression (Figure 5F-H).

Finally, for each unique lncRNA within a window of 5 kbp from the chosen OR, we examined the correlation of its expression level with that of upstream, chosen, and downstream ORs. Averaging across all unique lncRNAs confirmed that lncRNA expression is associated with a decrease in upstream OR expression and an increase in the expression of both the chosen OR and any downstream ORs (Figure 5I). This suggests that, as the promoter increases its transcriptional activity, the resulting increase in lncRNA expression serves to shut down the production of upstream transcripts.

These results help clarify why some cells in our heatmaps of tandem array expression show non-zero expression of ORs upstream of the chosen OR. For example, if we isolate all cells that express 9E121 as the chosen OR, we find that the heatmap becomes markedly cleaner when restricting the sample to cells with detectable coexpression of the upstream antisense lncRNA (Figure 5J) because these cells have greatly reduced expression of upstream ORs (Figure 5K).

To examine whether this pattern of lncRNA expression generalizes to ants and other insects, we analyzed published snRNA-seq data from the antennae of the Indian jumping ant *Harpegnathos saltator*^63^ (Figure S6A) and the honeybee *Apis mellifera*^37^ (Figure S6B). In both species, we identified tandem arrays that exhibit the staircase-like pattern of coexpression described in *O. biroi*, where the chosen OR is coexpressed with downstream ORs (Figures S6C and S6D). Publicly available genome annotations from *H. saltator*^64^ and *A. mellifera*^65^ contained twelve and eleven antisense lncRNAs nested within OR tandem arrays, respectively (Figures S6E and S6F). First, we plotted the coexpression of ORs neighboring lncRNAs and found that, as in *O. biroi*, lncRNA expression was associated with the expression of ORs immediately upstream (Figures S6G-J). We then analyzed the correlation of lncRNA expression with the expression of upstream, chosen and downstream ORs. As in *O. biroi*, lncRNA expression was associated with a decrease in upstream OR expression and an increase in downstream OR expression (Figures S6K and S6L). Taken together, our results suggest a comprehensive model of how transcriptional activity generates OR selectivity in hymenopterans and possibly other insects (Figure 6).

**Figure 6.**
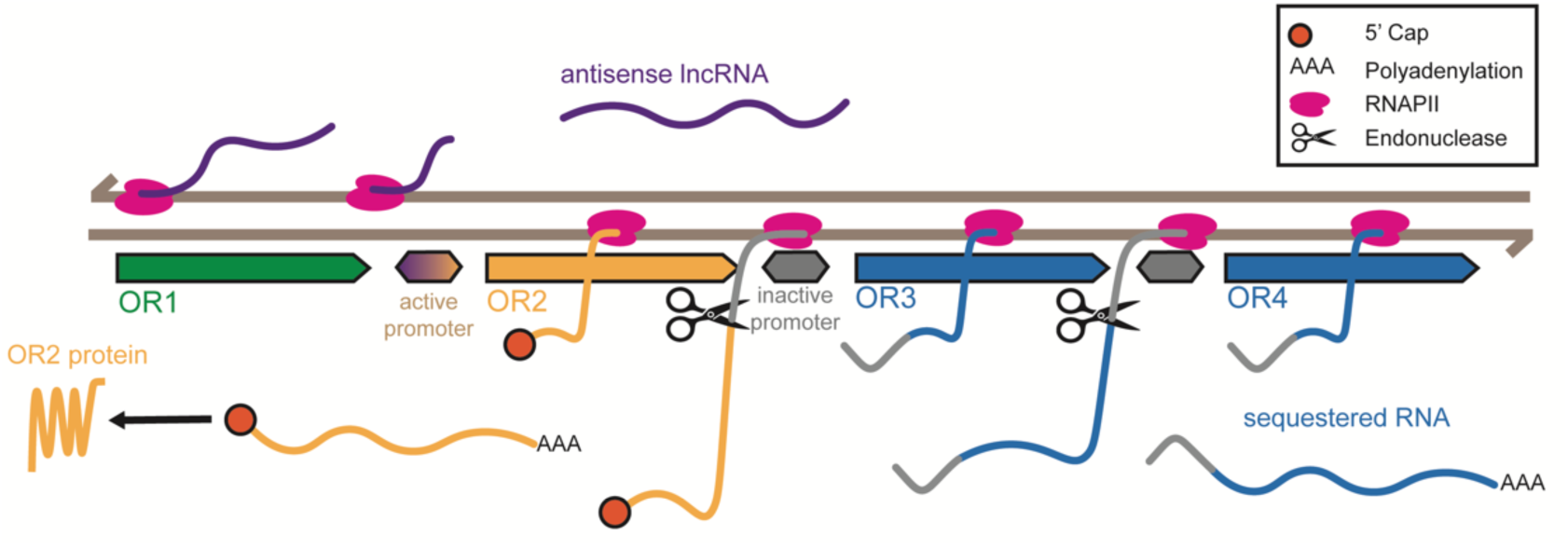
Non-coding Transcriptional Activity Enhances OR Selectivity at Tandem Arrays. Schematic model of transcriptional regulation at a hypothetical OR tandem array. A single bidirectional promoter initiates RNAPII activity in both directions. In the coding direction (left to right), RNAPII transcribes the chosen OR (yellow), and the transcript is capped at the 5′ end. Upon encountering the polyadenylation signal, the nascent transcript is cleaved and polyadenylated. RNAPII continues transcription past the 3′ end of the chosen OR into the intergenic region and beyond, producing downstream OR transcripts (blue) that are also polyadenylated but remain in the nucleus. We suspect that the intergenic region is retained in the 5’ UTR of each downstream gene. In the non-coding direction (right to left), RNAPII transcribes an antisense lncRNA (purple) that inhibits transcription of upstream ORs (green). Most of these lncRNAs are not polyadenylated. Only the capped transcripts from the chosen OR are exported into the cytoplasm and translated into functional protein.

### Bidirectional Transcription Ensures Monogenic Expression in the Case of Inversions

If correct, our model should also explain patterns of gene expression in the rare cases of OR inversions. While most tandem arrays are composed of genes oriented head-to-tail, we identified a few exceptions in which one or multiple genes were inverted relative to the rest of the tandem array. These genes provide a unique opportunity to test our model, particularly in cases where the antisense lncRNA spans the coding sequence of an upstream OR.

First, we examined T51, a tandem array of 16 genes in which 11 genes in the middle of the array are flipped in orientation (Figure 7A). 9E89, the last OR in the array, is coexpressed with all other ORs, which is expected because OSNs with a chosen OR in the same orientation should express it as a downstream OR, while the inverted genes 9E92-9E102 should express it as part of an antisense lncRNA. We co-stained 9E89 and 9E99 (Figure 7B) and found 9E89 reliably coexpressed in OSNs with 9E99 as the chosen OR, with nuclear transcript localization (Figure 7C and 7D). Similarly, OSNs with 9E89 as the chosen OR showed nuclear localization of 9E99 transcripts (Figure 7C and 7E).

**Figure 7.**
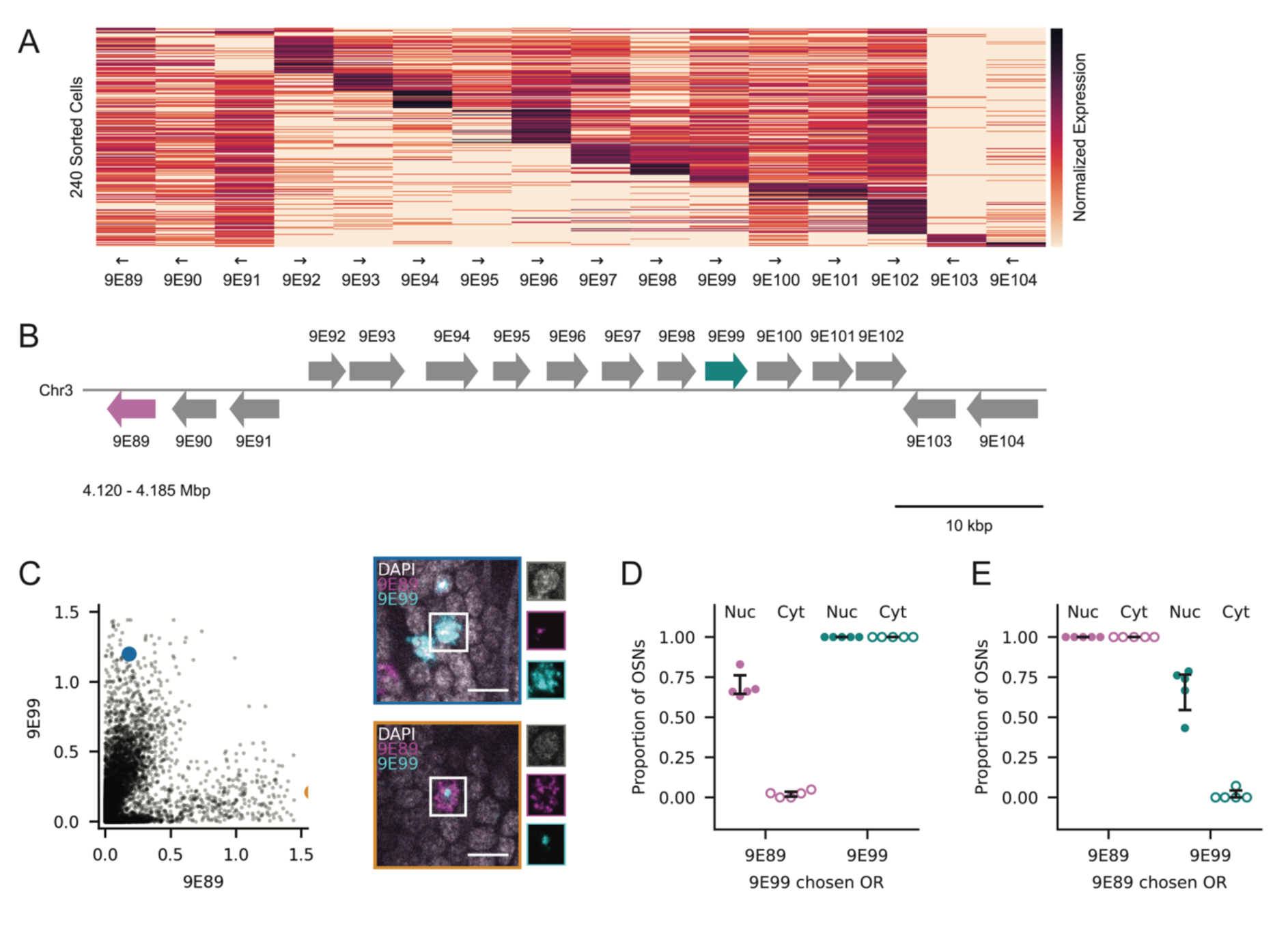
Non-coding Transcription Explains Expression Patterns at OR Gene Inversions. (A) Heatmap of log-normalized expression of all ORs in T51 across cells with a chosen OR in T51. Cells (rows) are sorted by the genomic position of their chosen OR. Arrows indicate strand orientation. (B) Schematic of T51, highlighting ORs 9E89 (magenta) and 9E99 (cyan). (C) Normalized nuclear signal for 9E99 and 9E89 in segmented OSN nuclei from n=5 antennae. Images with colored borders reflect the cells labeled with the corresponding colors and each channel is shown individually to the right of each image. Blue: cell with cytoplasmic 9E99 and nuclear 9E89. Orange: cell with cytoplasmic 9E89 and nuclear 9E99. Scale bars: 5 µm. (D-E) Proportion of OSNs with 9E99 (D) or 9E89 (E) as the chosen OR per antenna (n=5) with 9E89 and 9E99 signal in the nucleus and cytoplasm. Error bars: 95% CI centered on the mean.

Finally, we examined OR 9E198, which is located in tandem array T35 and flipped relative to all other ORs in that tandem array (Figure S7A). 9E198 is coexpressed with all ORs in the tandem array other than 9E200 and 9E201, the two ORs downstream of 9E198 (Figure S7A). We co-stained 9E196, 9E197 and 9E198 (Figure S7B) and observed that cells that had chosen 9E196 coexpressed the upstream gene 9E198 but not 9E197 (Figure S7C). Similarly, cells that had chosen 9E197 coexpressed both the downstream gene 9E196 and the upstream gene 9E198 (Figure S7D). In both cases, the non-chosen OR transcripts remained nuclear (Figure S7C and S7D).

These examples of inverted ORs show that ant OSNs reliably prevent ORs neighboring the chosen OR from producing protein products, regardless of the relative orientation of these genes. The mechanism of tandem gene duplication typically results in gene copies oriented in the same direction on the same strand.^33^ However, the mechanisms described here robustly ensure monogenic expression even when inversions do occur.

## DISCUSSION

The common tandem arrangement of chemosensory genes in insects^25–34^ and vertebrates^66,67^ presents a unique regulatory challenge. The accumulation of transcription factors at any single gene could lead to off-target transcription of nearby genes, and if these genes also produce protein, the tuning of the respective OSN may be affected. Here, we describe a novel solution to this problem that uses a bidirectional transcriptional interference mechanism. Increasing transcriptional activity on both sides of an active promoter in the form of downstream readthrough and upstream antisense lncRNAs enhances OR selectivity at the protein level.

This study builds on previous work in honeybees^37^ and mosquitoes^36^ showing that single promoters can drive expression of 2-6 OR genes arranged in tandem. In *O. biroi*, we extend this idea to tandem arrays containing dozens of genes, and we further propose that a novel form of transcriptional readthrough explains the data we observe. In our model, RNAPII continues transcribing after each cleavage event, producing independent polyadenylated transcripts of which only the first is exported out of the nucleus. We suspect that this transcriptional activity serves as a protective barrier that prevents the activation of downstream promoters. Our model differs from the mechanism governing a tandem array of three ionotropic receptor genes in *Drosophila*, where RNAPII produces a long polycistronic mRNA that spans all three receptors and lacks the first exons of the downstream genes, prohibiting their translation.^68^ While limited transcriptional readthrough (<5 kbp) typically occurs in healthy human cells,^69^ it appears that in ant OSNs, readthrough can extend significantly further from the active promoter (>100 kbp) to drive expression of non-translated OR transcripts.

The model we propose relies on an unusual failure of transcriptional termination that preserves proper cleavage. Canonically, once the nascent transcript is cleaved, RNAPII continues transcribing the downstream intergenic sequence until the exonuclease XRN2 degrades the leftover RNA fragment and dislodges RNAPII.^70–72^ Surprisingly, we find that each OR mRNA is cleaved and polyadenylated, yet RNAPII continues transcribing into the next OR gene, indicating a local barrier to termination. This unusual behavior is unprecedented at the scale we observe, although an example where a single RNAPII can produce two polyadenylated transcripts has been documented in vertebrates. Here, small nucleolar RNAs (snoRNAs) embedded immediately downstream of a PAS enable continued RNAPII transcription after cleavage.^73^ The snoRNAs recruit ribonucleoproteins co-transcriptionally, yielding a ribonucleoprotein cap that shields the freshly cleaved 5’ end from XRN2 and allows RNAPII to traverse multiple genes from a single promoter.^73^ An analogous protection mechanism could underlie OR-specific readthrough in ants, obviating the need to globally modulate XRN2 activity while ensuring that transcripts from downstream genes remain sequestered.

OR genes in the ant produce two qualitatively distinct mRNA species. The transcripts of chosen ORs are plentiful and exported into the cytoplasm, while transcripts produced via non-canonical readthrough are low in abundance and typically colocalize in the nucleus with the brightest signal from the chosen OR. We demonstrate that these downstream transcripts are spliced normally, and hypothesize that the distinguishing feature of chosen OR transcripts is the presence of a 5’ cap, a 7-methylguanosine moiety attached to the first transcribed nucleotide.^41–43^ In the absence of this cap, downstream transcripts would be vulnerable to post-transcriptional degradation. Although downstream genes may be transcribed at similar rates to the chosen OR, increased degradation could account for their low steady-state levels.

We further propose that ant ORs possess bidirectional promoter regions that generate antisense lncRNAs. Although bidirectional gene pairs are common in insects,^74^ promoters in *Drosophila* are predominantly unidirectional.^75^ In the human genome, most promoters are strongly directional when assessing mature transcripts, but many appear bidirectional when measuring nascent transcripts.^62^ We present evidence that bidirectional initiation is common in the ant genome, but antisense elongation is specific to OR promoters.

Non-coding transcripts have emerged as important mediators of transcriptional interference and gene silencing,^76,77^ and the abundance of lncRNAs in the *Drosophila* antenna^56^ has led to speculation on the involvement of lncRNAs in insect OSNs.^57^ In some cases, lncRNAs themselves are critical for silencing,^78^ whereas in others, the act of transcription alone is sufficient to repress gene expression.^79^ Transcriptional interference is thought to occur through several mechanisms, including promoter occlusion and RNAPII collision with transcription factors.^79^ RNAPII can engage in transcriptional interference irrespective of whether it transcribes in the same^68,76,80–83^ or opposite^84,85^ orientation as the gene it is silencing. However, all studies to date on lncRNA-associated interference in tandemly arrayed genes have focused on gene pairs or small clusters,^68,76,86^ rather than large tandem arrays containing dozens of genes. Our analysis suggests that sense and antisense transcripts from a bidirectional promoter region can silence dozens of genes located up to >100 kbp upstream or downstream. We suspect that neither the sense nor the antisense transcripts serve a direct functional role beyond transcriptional interference, preventing initiation from nearby promoters.

Social insects rely extensively on chemical communication, and the rapid evolutionary turnover of ORs, particularly in ants, is believed to underpin between-species differences in behaviors such as non-nestmate discrimination and prey recognition, as well as various collective activities.^6,87–91^ Here, we present evidence suggesting that bidirectional transcription from OR promoter regions ensures OR selectivity in dense, gene-rich tandem arrays. Remarkably, this mechanism is active even at singleton OR genes and OR genes that are inverted within their respective tandem array, showing that it emanates from the promoter region itself. This opens the possibility that when ant ORs are duplicated together with this conserved regulatory region, they immediately give rise to a novel type of OSN that exclusively produces the respective receptor. This transcription-based mechanism might thus be crucial to maintaining monogenic OR selectivity in a clade characterized by frequent gene duplication events.

## RESOURCE AVAILABILITY

### Lead Contact

Further information and requests for resources and reagents should be directed to and will be fulfilled by the lead contact, Daniel J. C. Kronauer.

### Materials Availability

All materials other than ants are commercially available, and ants can be provided upon request in accordance with federal regulations.

### Data and Code Availability

Confocal RNA-FISH images are publicly available via the Brain Image Library. Links and accession numbers are provided in the Key Resources Table. All code used for image segmentation, alignment, analysis, quantification, and figure generation can be found on GitHub (https://github.com/Social-Evolution-and-Behavior/Glotzer-Kronauer-2025).

## ACKNOWLEDGEMENTS

We thank Bayley McDonald and Sascha Duttke for acquiring and sharing the csRNA-seq data. Stephany Valdés-Rodríguez, Alek Rahman and Alejandra Hurtado-Giraldo maintained stock colonies of ants used for experiments. We thank Kip Lacy for his advice on quantifying the rRNA-sequencing data, Anindita Brahma for her input on the manuscript, Shixin Liu, Bob Darnell and Vanessa Ruta for valuable discussions, and other members of the Kronauer lab for their feedback. This is Clonal Raider Ant Project paper number 38.

## AUTHOR CONTRIBUTIONS

G.L.G. analyzed the sequencing data, designed RNA-FISH probes, conducted RNA-FISH experiments, and wrote code used for segmentation, analysis, and figure generation. P.D.H.P. provided input on the planning of experiments, design of RNA-FISH probes, and data analysis. G.L.G. and D.J.C.K. designed the experiments and wrote the paper, and all authors provided feedback on the manuscript. D.J.C.K. supervised the project.

## METHODS

### KEY RESOURCES TABLE

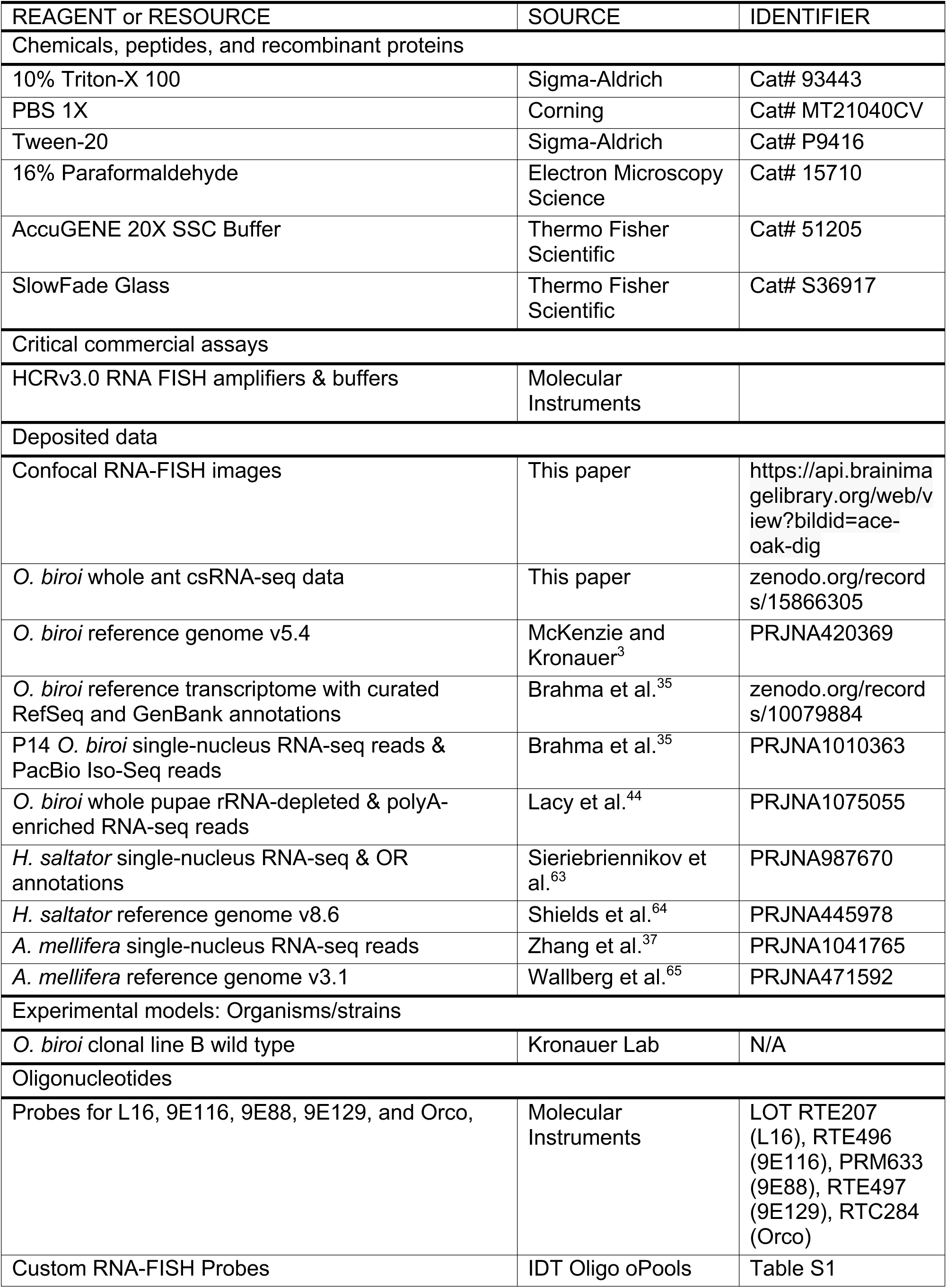
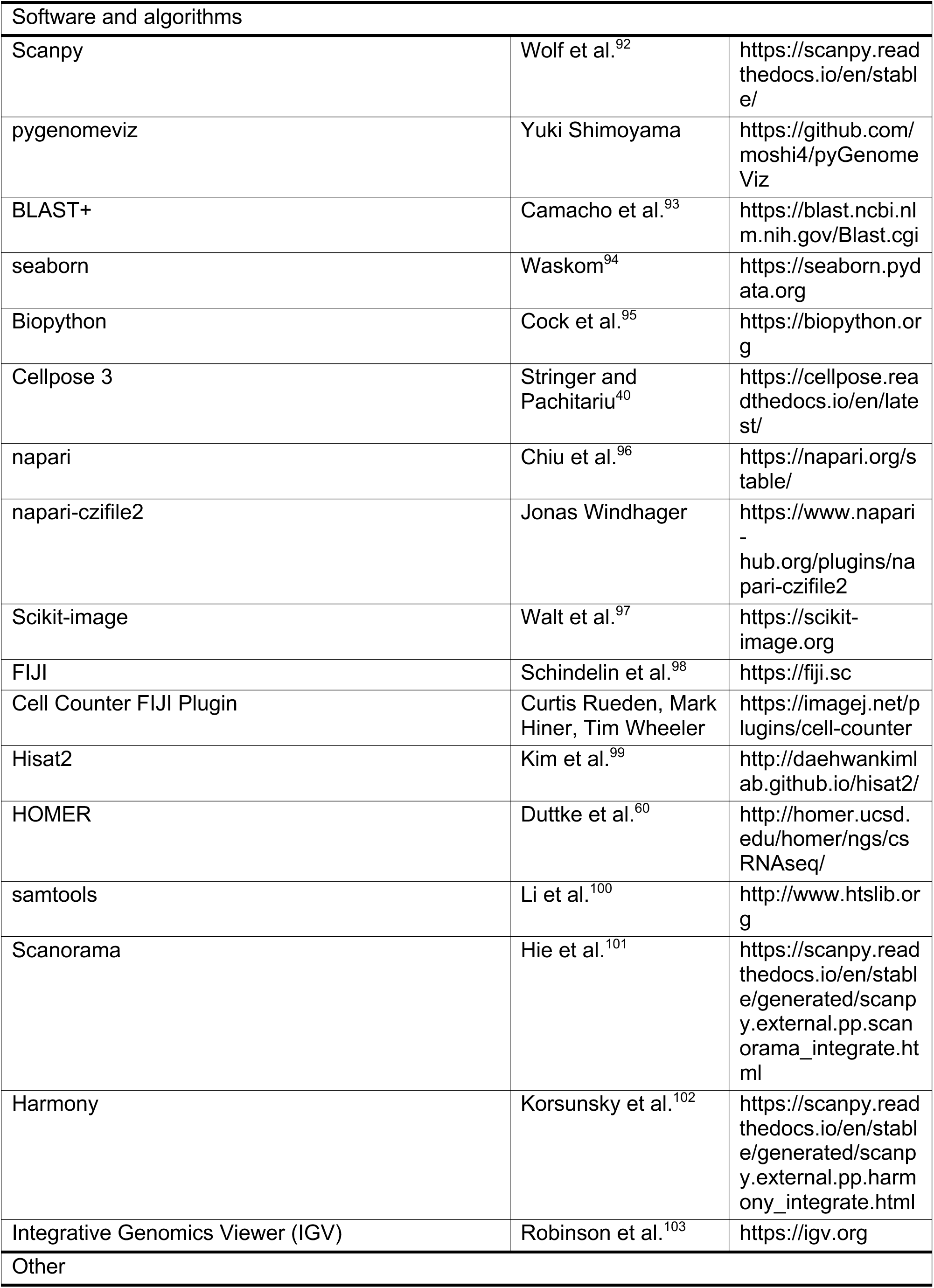
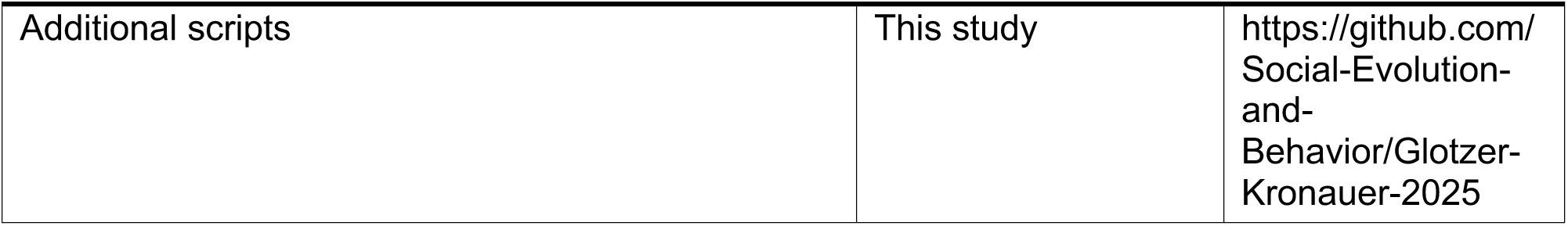

### EXPERIMENTAL MODEL AND STUDY PARTICIPANT DETAILS

#### Ant Husbandry and Maintenance

*Ooceraea biroi* ants were obtained from large stock colonies maintained at 25 °C in Tupperware containers (32x14 cm) with a plaster of Paris floor. During the brood care phase, colonies were fed three times per week with frozen *Solenopsis invicta* (fire ant) brood and cleaned and watered weekly as needed. Pupae from clonal line B were collected on the day of pupation (P0) and housed with adult workers in small Petri dishes also lined with plaster of Paris. Pupae were aged to 14 days (P14), at which point antennal clubs were dissected.

### METHOD DETAILS

#### rRNA-depleted and polyA-enriched RNA-seq Data

To assess RNA sequencing coverage of OR and non-OR gene pairs, we used existing datasets of rRNA-depleted and polyA-enriched RNA-sequencing (PRJNA1075055) from whole *O. biroi* pupae.^44^ We retrieved the curated gene annotations that include RefSeq (GCF_003672135.1) and GenBank (GCA_003672135.1) annotations in GTF format from Zenodo (https://zenodo.org/records/10079884).

For our analysis of gene pairs, we first filtered for genes with a coding sequence length between 100 bp and 10 kbp. Candidate gene pairs were selected based on an intergenic distance ranging from 50 bp to 10 kbp and were required to be located on the same strand. Pairs were excluded if either gene or the intergenic region overlapped with any other annotated gene on either strand. This filtering yielded 211 OR gene pairs, and we randomly sampled an equal number of non-OR gene pairs for comparison.

Stranded base-pair read coverage was calculated directly from raw BAM files using Samtools.^100^ For each upstream gene in each pair, we calculated the mean coverage across all annotated exons and introns. For intergenic regions, mean coverage was calculated across the entire intergenic span. To account for expression differences, relative coverage was computed by normalizing to the mean exon coverage of the upstream gene in each pair.

#### csRNA-seq Data

To assess the exact location of transcriptional start sites (TSSs), we analyzed an unpublished dataset of capped short RNA-sequencing (csRNA-seq) from whole adult *O. biroi* ants. The data was acquired by Bayley McDonald, a graduate student in the lab of Dr. Sascha Duttke at Washington State University, following an established protocol.^60,61^ Bulk extracted RNA was size selected and enriched for 5’-capped RNAs. After amplification, strand-specific paired-end libraries were depleted of rRNA and sequenced on an Illumina NextSeq 2000. Sequencing reads were trimmed using HOMER^60^ and aligned to the *O. biroi* reference genome v5.4 using Hisat2.^99^ A multimapping threshold of 10 was used to ensure unique alignment. The resulting bedGraph files were loaded into Python for subsequent analysis.

Many csRNA-seq peaks were within the annotated UTR of an OR gene, indicating that some of our gene models have incorrect TSSs. Thus, we assigned csRNA-seq reads to OR genes by searching the 1 kbp upstream of the first CDS. For the analysis of non-ORs, the same 211 gene pairs were used as in the rRNA-depleted RNA-seq analysis. This sampling ensured that the downstream gene TSS was not in close proximity to an existing antisense gene annotation. Here, we searched the 1 kbp upstream of the first CDS for ORs and the 1 kbp upstream of the annotated TSS for non-ORs.

#### Long-read RNA-seq Data

To produce the image of long-read sequencing coverage of T79 in Figure S2D, we loaded the BAM files of aligned long-read Isoseq reads from P14 antennae (PRJNA1010363)^35^ into Integrative Genomics Viewer^103^ and exported an image of the locus containing T79.

#### Ooceraea biroi snRNA-seq Data

We used a published snRNA-seq dataset from wild-type P14 *O. biroi* antennae.^35^ We specifically isolated only the neurons by thresholding for expression of previously-identified neuron markers.^35^ Each nucleus had at least two of the five markers (LOC105284916, LOC105280759, LOC105285306, LOC105276401, LOC105275115) with a UMI ≥2.

#### Harpegnathos saltator snRNA-seq Data

We analyzed three published snRNA-seq datasets from wild-type adult *H. saltator* antennae^63^. Raw gene expression matrices were processed using Scanpy,^92^ and datasets were integrated with batch correction using Scanorama.^101^ OR and lncRNA genes were identified using the *H. saltator* reference genome v8.6.^64^ OR genes were assigned to tandem arrays using cluster definitions from the supplementary materials of Sieriebriennikov et al.^63^

#### Apis mellifera snRNA-seq Data

We analyzed three published snRNA-seq datasets from the antennae of wild-type forager, nurse, and newly emerged *A. mellifera*.^37^ We integrated the datasets using Harmony.^102^ OR gene annotations were obtained from the supplementary materials of Zhang et al.^37^ Tandem arrays were defined as clusters of non-overlapping ORs separated by ≤10 kb. lncRNAs were annotated using the *A. mellifera* reference genome v3.1.^65^

#### Preprocessing of snRNA-seq Data

Each dataset was imported into Scanpy^92^ and gene expression values were log-transformed and normalized to a target sum of 10,000 counts per cell.

#### Quantification of lncRNAs using snRNA-seq Data

To identify nested lncRNAs, we searched for transcripts overlapping in genomic coordinates with OR tandem arrays. For each nested lncRNA, we annotated the nearest upstream and downstream OR genes within the same array, relative to the lncRNA TSS. We calculated the coexpression of each OR gene in all cells expressing the corresponding lncRNA (>0 counts). Genomic distance between each lncRNA and OR gene was defined as the distance between their TSSs.

To assess expression correlation, we identified cells in which the chosen OR (the highest-expressing OR per cell) was located within 5 kbp upstream of an antisense lncRNA. In these cases, distance was calculated between the OR TSS and the lncRNA 3’ end.

#### Sample Preparation for RNA-FISH

Line B ants were aged to 14 days post-pupation (P14). Ants were washed for 30 seconds in ice-cold 95% ethanol followed by 1xPBS. Antennae were dissected in PBS using microdissection scissors and transferred to 1.5 mL microcentrifuge tubes containing 1 mL of 4% paraformaldehyde (PFA) in 1xPBS with 0.5% Triton X-100. The samples were incubated in the 4% PFA solution for one hour at room temperature (RT) on a rocker. Antennae were then sonicated on a cooling block using a Q700 Sonicator (QSonica, Newtown, CT) fitted with a 1.6mm microtip. Sonication was performed at amplitude 15 with the following parameters: 20 cycles of 2 seconds on and 20 seconds off. Following sonication, samples were returned to RT and fixed for an additional hour on a rocker. After fixation, the antennae were washed in 0.1% PBS-Tween (PBST) for 5 minutes and dehydrated using an ice-cold methanol/PBST gradient (25%, 50%, 75%, 100%) for 10 minutes at each step. Samples were then bleached in 3% hydrogen peroxide (H₂O₂) in methanol for 1 hour at 4 °C under bright light. Antennae were returned to 100% methanol and stored at -20 °C until staining. Prior to staining, samples were rehydrated stepwise using the reverse methanol/PBST gradient (75%, 50%, 25%, 0%).

#### RNA-FISH Probes

For the genes L16, 9E116, and 9E88, we used probes designed by Molecular Instruments (Los Angeles, CA). For all other genes, tandem arrays, intergenic regions, and putative lncRNAs, we designed custom RNA-FISH probes compatible with the Molecular Instruments amplification system. To ensure probe specificity, we used command-line BLASTN^93^ against the *O. biroi* transcriptome and excluded any target regions that shared consensus sequences with other genes. For intergenic and putative lncRNA probes, we further excluded regions that were not unique to each intergenic and putative lncRNA. We designed 30 probe pairs against each target, unless the unique regions were not long enough, in which case we ordered the maximum number of probes that would fit. All probe design code is available on GitHub (https://github.com/Social-Evolution-and-Behavior/Glotzer-Kronauer-2025), and a complete list of custom probe sequences is provided in Table S1. Oligonucleotide pools were synthesized by IDT as oPools (50 pmol per oligo) and reconstituted in 50 µL of nuclease-free water to generate a 1 µM stock solution.

#### RNA-FISH Staining

We followed the HCR v3.0 FISH protocol for chicken embryos as described by Choi et al.,^104^ with slight modifications. Hybridization was carried out using 4 µL of 1 µM probe stock in 300 µL of probe hybridization buffer at 37°C for 16 hours. Amplification was performed for 16 hours in the dark using 6 µL of each hairpin in 300 µL of amplification buffer, with 1 µL DAPI added. Samples were mounted at room temperature in SlowFade Glass mounting medium (Thermo Fisher, Cat# S36917).

#### Confocal Microscopy

Antennae were imaged on a Zeiss LSM 900 confocal microscope using 405 nm, 488 nm, 561 nm, and 633 nm lasers. We used a Zeiss LD LCI Plan-Apochromat 40X / 1.2NA multi-immersion objective lens immersed in glycerol to acquire z-stack images at 0.8x zoom with a z-step of 1 µm, capturing slices at 2048x2048 pixel resolution. Laser power was calibrated individually for each stain to avoid underexposure or saturation. For samples stained in parallel, identical laser settings were used to enable accurate and consistent quantification of RNA-FISH signal.

#### Cell Segmentation of OSN Nuclei

Confocal images were loaded using the napari-czifile2 plugin (www.napari-hub.org/plugins/napari-czifile2). We trained a custom 2D nuclei segmentation model for OSNs by fine-tuning the default Cellpose 3 nuclei model.^40^ We hand-labeled 24 representative z-slices extracted from six images to use as ground truth, with 16 slices used for training and 8 for testing. We specifically only labeled OSNs and did not label support cells, which are elongated, or IR-expressing cells, which have larger nuclei. The model was trained for 500 epochs with a learning rate of 0.005 and weight decay of 10^-4^, using an estimated nuclear diameter of 3 µm. We monitored training and validation loss curves and confirmed convergence of the test loss to approximately 0.2. The model was further validated on held-out images to ensure accurate segmentation. Following validation, the trained model was applied to all images, and the resulting nuclear regions of interest (ROIs) were saved for downstream analysis.

#### Quantification of Nuclear and Cytoplasmic Intensity

Each image and its corresponding nuclear ROIs were processed to quantify signal intensities. To correct for depth-dependent signal attenuation, we normalized the intensity of each non-DAPI channel across z-slices using the mean DAPI intensity as a reference. Specifically, we scaled each slice using a scalar defined as the ratio of the maximum DAPI intensity across slices to the DAPI intensity of the current slice.

Background subtraction was performed on each z-slice using a Gaussian filter with a sigma of 100 and FISH signal was segmented by applying the Triangle threshold from scikit-image.^97^ The signal mask was further improved by removing small objects (<12 pixels in area). For each nuclear ROI, we recorded the mean signal intensity across channels, along with centroid coordinates and area.

To quantify cytoplasmic signal, each nuclear ROI was dilated by 3 pixels (∼270 nm), reflecting the estimated cytoplasmic thickness in OSNs. The cytoplasmic region was defined using a binary subtraction of the original nucleus from the dilated mask, preserving the cytoplasmic periphery. Overlapping regions, either between cytoplasmic masks or between cytoplasmic and neighboring nuclear masks, were excluded to avoid signal contamination in dense regions. For each cytoplasmic ROI, we quantified the mean signal intensity in all channels.

#### Normalization of Nuclear and Cytoplasmic Signal

To account for inter-image variability and differences in imaging conditions, nuclear and cytoplasmic mean signal intensities were normalized independently for each image and each channel using robust quantile scaling. For each distribution, the lower (0.001) and upper (0.999) quantiles of the nuclear signal defined the normalization range, effectively reducing the impact of outliers. The same nuclear quantile range was used to normalize the cytoplasmic signal.

#### Labeling of Chosen OR-Expressing Cells

For all RNA-FISH experiments, we designated cells as expressing an OR as the chosen OR if it expressed an OR with a normalized nuclear signal >0.75 and a normalized cytoplasmic signal >0.2. Additionally, we applied standard size selection, ensuring that these cells had a nuclear area between 400-900 pixels (3.2-7.2 µm^2^) and cytoplasmic area >100 pixels (>0.8 µm^2^). We further selected for circular nuclei by ensuring that the ROIs had an eccentricity <0.8. The number of segmented cells in each antenna may be slightly different than the counts provided in this paper because the z-step we used (1 µm) is small enough that the same cell may be counted twice in two different z-planes. However, this does not systematically affect our results, as we are interested in the proportion of cells that coexpress two genes, rather than their raw counts.

#### Quantification of Nuclear and Cytoplasmic Colocalization

A cell was classified as exhibiting nuclear transcript localization if its normalized nuclear signal was >0.1. Cells were further classified as exhibiting cytoplasmic transcript localization if they also had a normalized cytoplasmic signal >0.2. These thresholds were uniformly applied across all channels, images, and experimental replicates to ensure consistency.

#### Quantification of Subnuclear RNA-FISH Expression Domains

RNA-FISH signal was segmented using the same method applied for signal intensity measurements. To quantify whether probes label the same RNA molecules, we measured the area of each signal domain and the area composed of the overlap between probes in different channels.

### QUANTIFICATION AND STATISTICAL ANALYSES

Analyses were performed in Python using standard scientific libraries including NumPy, Pandas, Seaborn and Matplotlib. Code used for analysis, quantification and plotting is available on GitHub (https://github.com/Social-Evolution-and-Behavior/Glotzer-Kronauer-2025). Statistical tests and p-values are noted in the figure legends.

## SUPPLEMENTAL VIDEO AND EXCEL TABLE TITLES AND LEGENDS

**Supplemental Video 1. Nuclear Segmentation of OSNs**

Example of nuclei segmentation applied to each confocal z-stack for use in RNA-FISH analysis. Scale bar: 50 µm.

**Supplemental Video 2. Cytoplasmic Segmentation of OSNs**

Example of cytoplasmic segmentation applied to each confocal z-stack for use in RNA-FISH analysis. Scale bar: 50 µm.

**Supplemental Table 1. Custom RNA-FISH Probe Sequences**

Excel file containing a sheet with the custom probe sequences used for RNA-FISH for each probe set.

---

**Figure S1.**
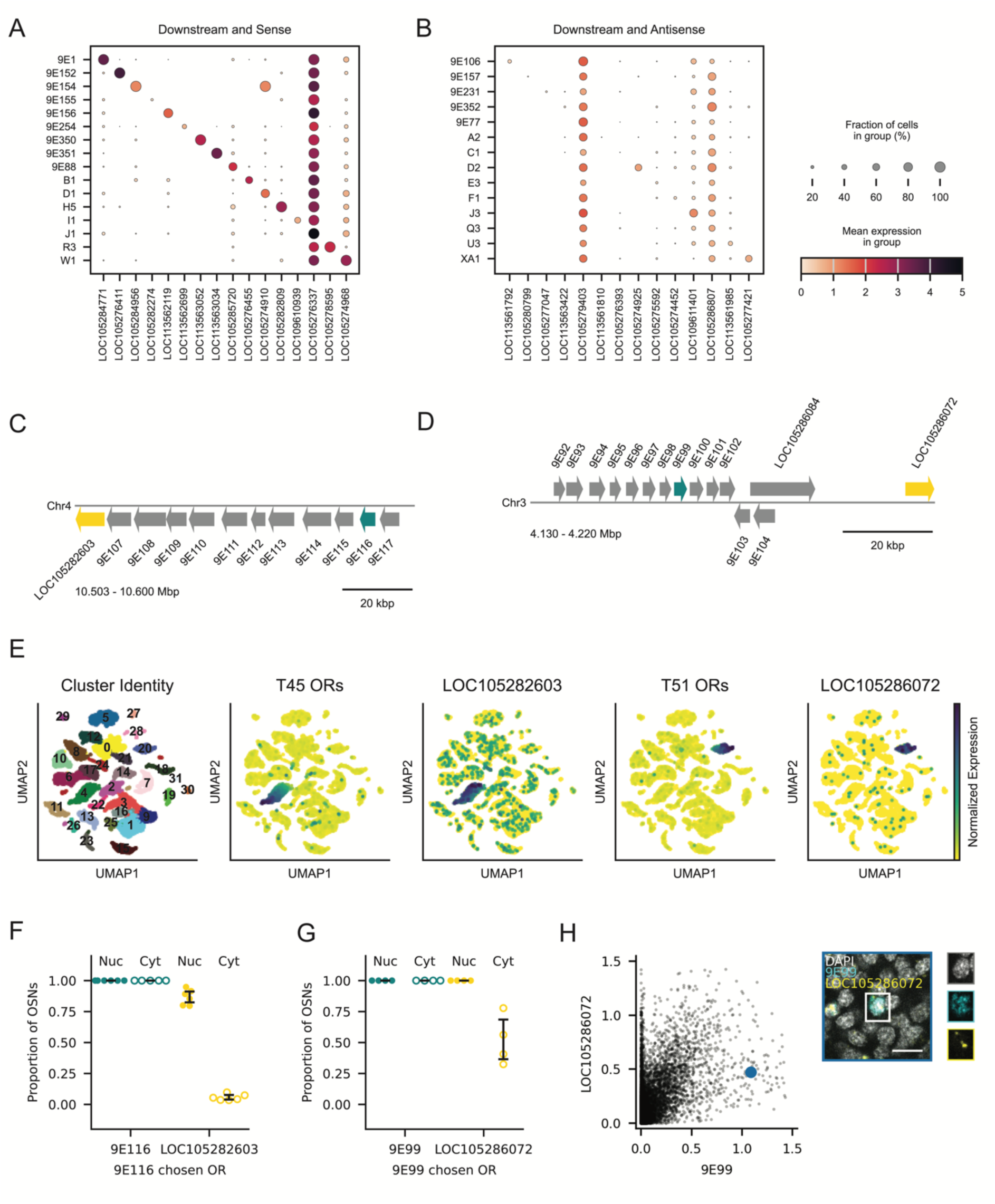
Additional Characterization of Non-OR genes, Related to Figure 1. (A-B) Expression of non-OR genes downstream of singleton ORs on the same strand (A) or opposite strand (B) as the focal OR. Column *n* corresponds to the downstream gene of the singleton OR in row *n*. Each group is composed of cells that express the OR as the chosen OR. Dot size corresponds to percentage of cells in each group that express a gene at a detectable level (>0) and dot color reflects the log-normalized expression level. (C) Schematic of a subset of T45 highlighting 9E116 (cyan) and LOC105282603 (yellow). 9E116 is located 81 kbp upstream of LOC105282603. (D) Schematic of a subset of T51 highlighting 9E99 (cyan) and LOC105286072 (yellow). 9E99 is located 51 kbp upstream of LOC105286072. (E) UMAPs of antennal neurons colored by cluster (left), mean expression of T45 ORs (second from left), expression of LOC105282603 (middle), mean expression of T51 ORs (second from right), expression of LOC105286072 (right). (F) Proportion of OSNs with 9E116 as the chosen OR per antenna (n=6) exhibiting 9E116 and LOC105282603 signal in the nucleus and cytoplasm. (G) Proportion of OSNs with 9E99 as the chosen OR per antenna (n=4) exhibiting 9E99 and LOC105286072 signal in the nucleus and cytoplasm. Error bars: 95% CI centered on the mean (F, G). (H) Normalized nuclear signal for LOC105286072 vs. 9E99 in segmented OSN nuclei from n=4 antennae. The image with a blue border shows an example cell with nuclear 9E99 and cytoplasmic LOC105286072 that is labeled in blue in the plot. Each channel is shown individually to the right. Scale bar: 5 µm.

**Figure S2.**
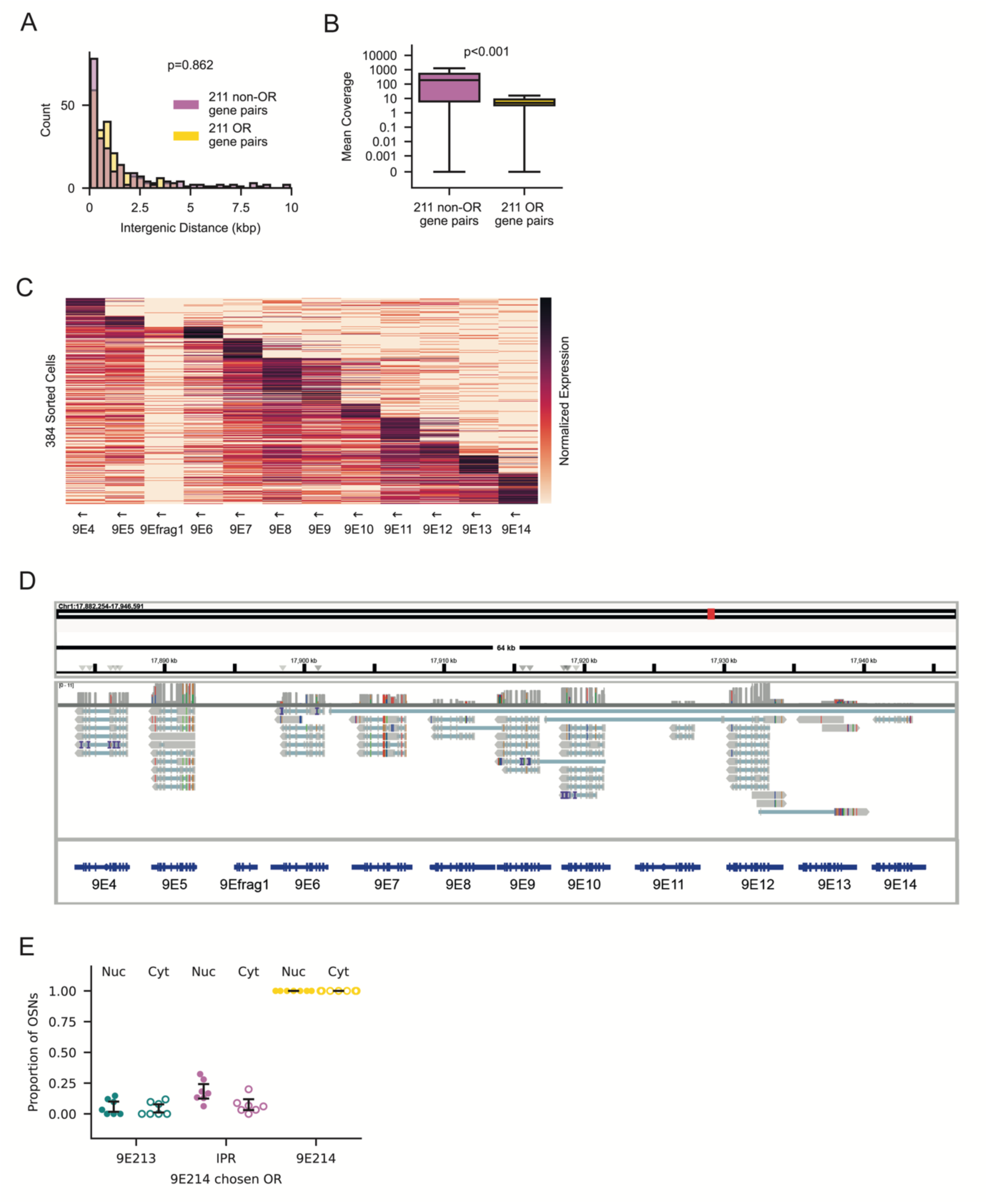
Additional Analysis of Intergenic Regions, Related to Figure 2. (A) Histogram of intergenic distances for 211 non-OR gene pairs (magenta) and 211 OR gene pairs (yellow). P-value from Wilcoxon rank-sum test. (B) Mean rRNA-depleted RNA-seq coverage across exons of 211 non-OR (magenta) and 211 OR (yellow) gene pairs. P-value from Wilcoxon rank-sum test. (C) Heatmap of log-normalized expression of all ORs in T79 across cells with a chosen OR in T79. Cells (rows) are sorted by the genomic position of their chosen OR. Arrows indicate strand orientation. (D) Alignment of long-read mRNA sequencing to the T79 locus. (E) Proportion of OSNs with 9E214 as the chosen OR per antenna (n=5) exhibiting 9E213, intergenic PR and 9E214 signal in the nucleus and cytoplasm. Error bars: 95% CI centered on the mean.

**Figure S3.**
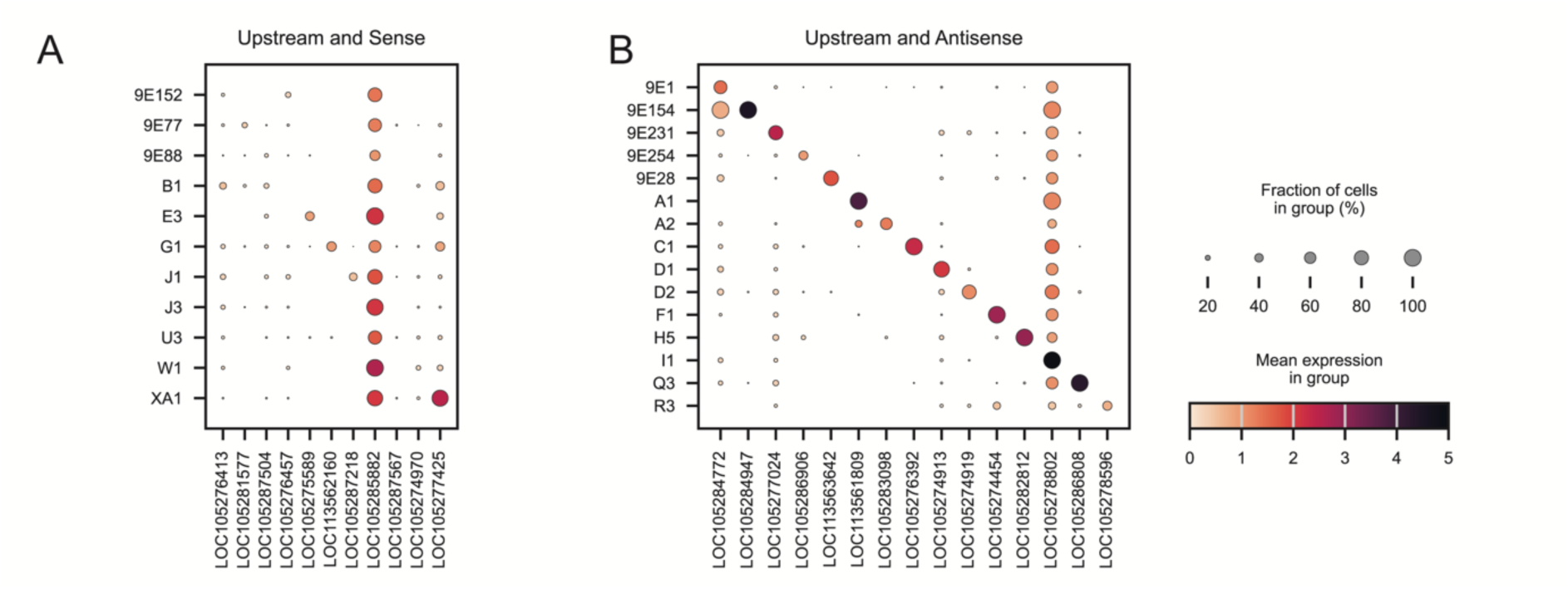
Characterization of Non-ORs Upstream of Singleton ORs, Related to Figure 3. (A-B) Expression of non-OR genes upstream of singleton ORs on the same strand (A) or opposite strand (B) as the focal OR. Column *n* corresponds to the upstream gene of the singleton OR in row *n*. Each group is composed of cells that express the OR as the chosen OR. Dot size corresponds to percentage of cells in each group that express a gene at a detectable level (>0) and dot color reflects the log-normalized expression level.

**Figure S4.**
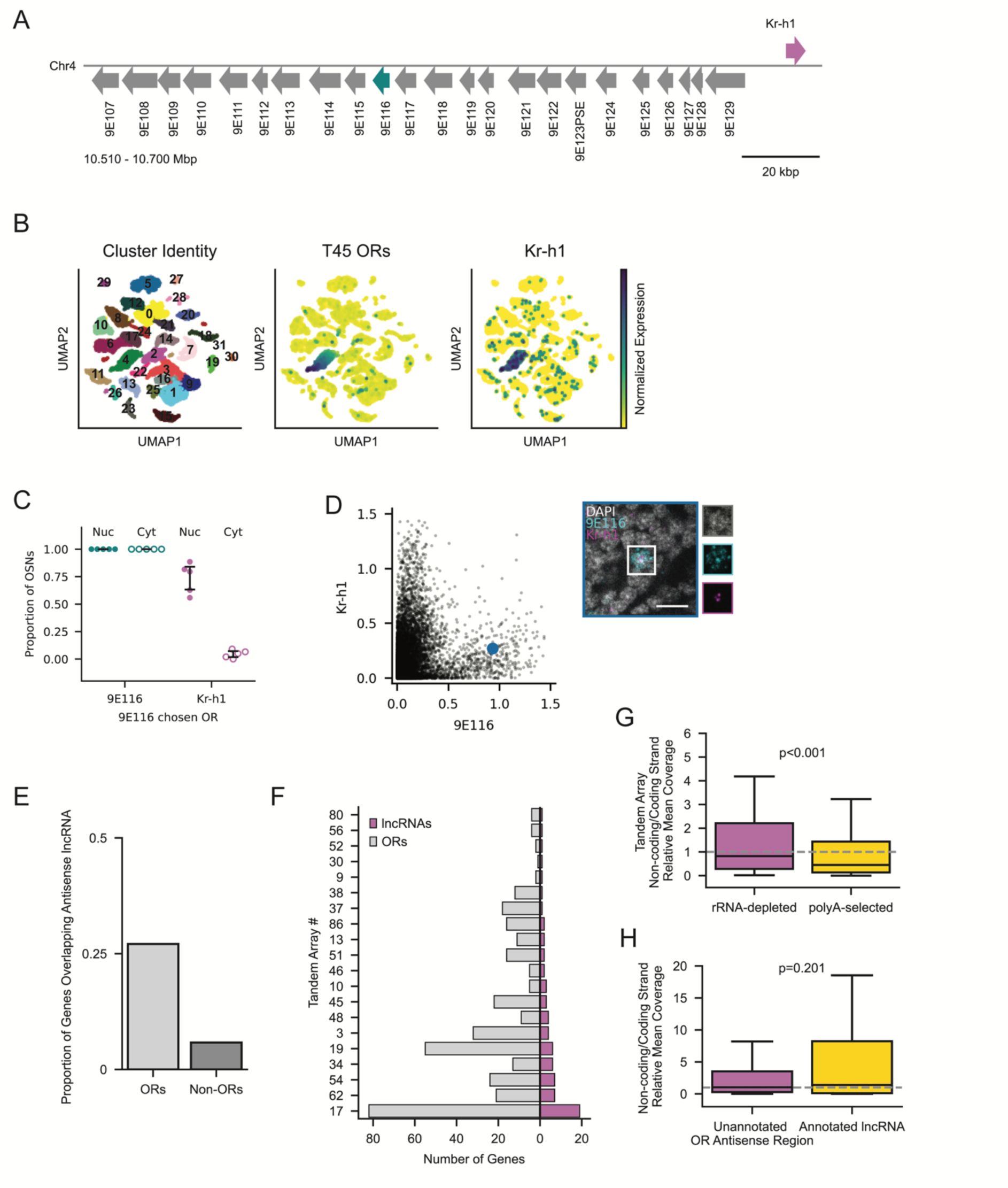
RNA from Upstream Antisense Non-OR Genes is Sequestered, Related to Figure 3. (A) Schematic of T45 highlighting 9E116 (cyan) and Kr-h1 (magenta). 9E116 is located 106 kbp upstream of Kr-h1. (B) UMAPs of antennal neurons colored by cluster (left), mean expression of T45 ORs (middle), and expression of Kr-h1 (right). (C) Proportion of OSNs with 9E1116 as the chosen OR per antenna (n=5) exhibiting 9E116 and Kr-h1 signal in the nucleus and cytoplasm. Error bars: 95% CI centered on the mean. (D) Normalized nuclear signal for Kr-h1 vs. 9E116 in segmented OSN nuclei from n=5 antennae. The image with blue borders shows an example cell with cytoplasmic 9E116 and nuclear Kr-h1 that is labeled in blue in the plot. Each channel is shown individually to the right. Scale bar: 5 µm. (E) Proportion of OR and non-OR genes that overlap with annotated antisense lncRNAs. (F) Number of antisense lncRNAs nested within each OR tandem array (magenta), and the number of OR genes per array (grey). (G) Ratio of non-coding to coding strand coverage for tandem arrays with ≥2 OR genes, using rRNA-depleted RNA-seq (magenta) and polyA-enriched RNA-seq (yellow). (H) Relative coverage of OR antisense regions (magenta) and annotated antisense lncRNAs nested in tandem arrays (yellow). Antisense coverage is normalized to the coding-strand coverage. (G, H) Each boxplot shows the median and quartiles; the whiskers extend to 1.5 times the interquartile range. P-value from Wilcoxon rank-sum test. Dotted line at y=1.

**Figure S5.**
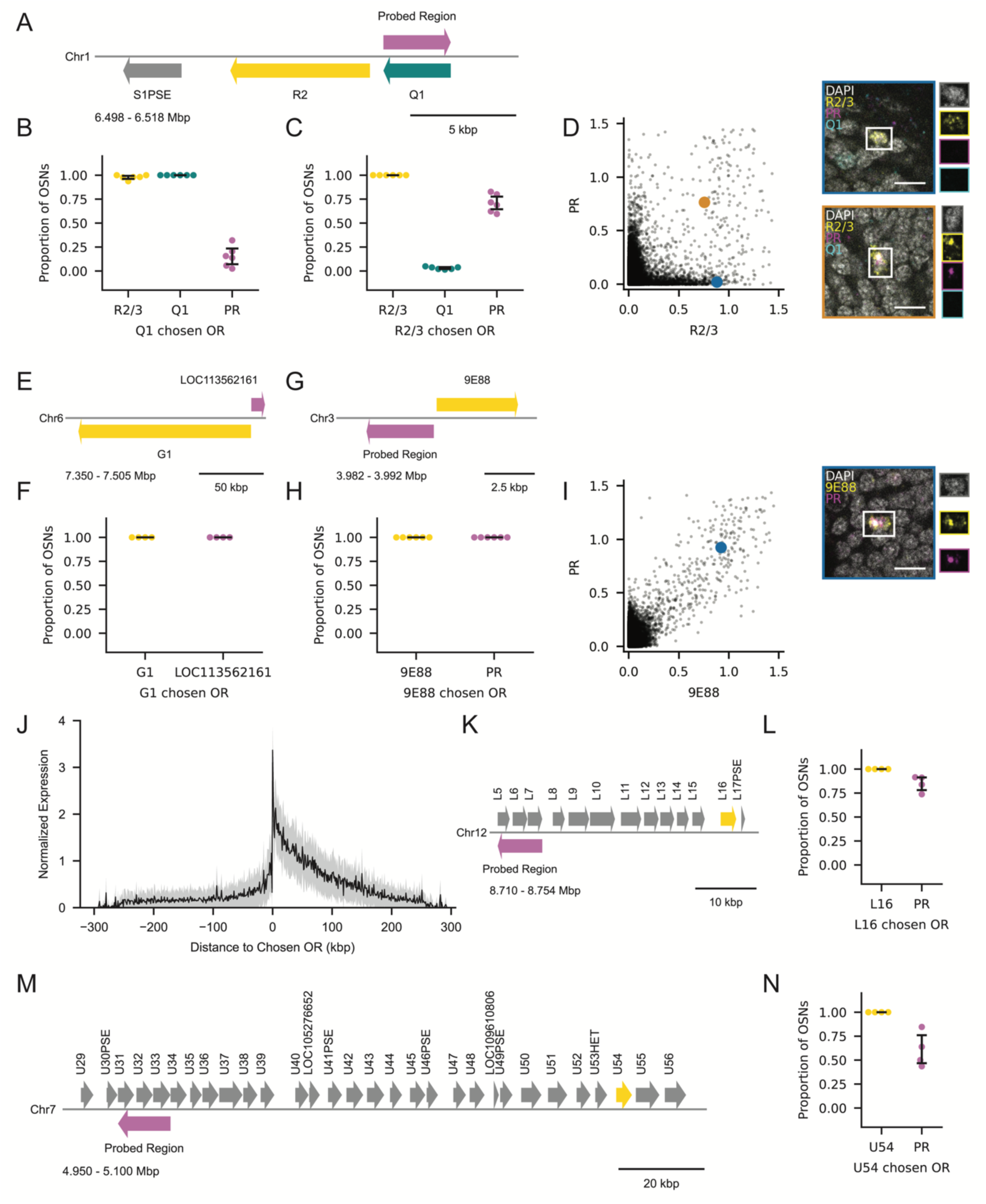
Additional Staining of lncRNAs, Related to Figure 4. (A) Schematic of T70, highlighting Q1 (cyan), R2 (yellow), and the probed region (PR) targeting a putative antisense lncRNA (magenta). (B-C) Proportion of OSNs with Q1 (B) or R2/3 (C) as the chosen OR per antenna (n=6) exhibiting Q1, R2/3 and PR signal in the nucleus. (D) Normalized nuclear signal for PR vs. R2/3 in segmented OSN nuclei from n=6 antennae. Images with colored borders reflect the cells labeled with the corresponding colors and each channel is shown individually to the right of each image. Blue: cell with only cytoplasmic R2/3. Orange: cell with cytoplasmic R2/3 and nuclear PR. (E) Schematic of the singleton OR G1 (yellow) and the antisense lncRNA LOC113562161 (magenta). G1 is 48 kbp away from the nearest other OR. (F) Proportion of OSNs with G1 as the chosen OR per antenna (n=4) exhibiting G1 and LOC113562161 signal in the nucleus. (G) Schematic of 9E88 and a probed region (PR) targeting a putative antisense lncRNA (magenta). (H) Proportion of OSNs with 9E88 as the chosen OR per antenna (n=6) exhibiting 9E88 and PR signal in the nucleus. (I) Normalized nuclear signal for 9E88 vs. PR in segmented OSN nuclei from n=6 antennae. The blue dot indicates an example cell with cytoplasmic 9E88 and nuclear PR. The cell is shown in the blue-ordered image, and each channel is shown individually to the right. (J) Mean and standard deviation of OR expression vs. genomic distance from the chosen OR TSS using snRNA-seq data. (K) Schematic of a subset of T3, highlighting L16 (yellow) and the probed region (PR) targeting a putative antisense lncRNA (magenta) 30 kbp upstream. (L) Proportion of OSNs with L16 as the chosen OR per antenna (n=4) exhibiting L16 and PR signal in the nucleus. (M) Schematic of a subset of T19, highlighting U54 (yellow) and the probed region (PR) targeting a putative antisense lncRNA (magenta) 103 kbp upstream. (N) Proportion of OSNs with U54 as the chosen OR per antenna (n=4) exhibiting U54 and PR signal in the nucleus. Error bars: 95% CI centered on the mean. Scale bars: 5 µm.

**Figure S6.**
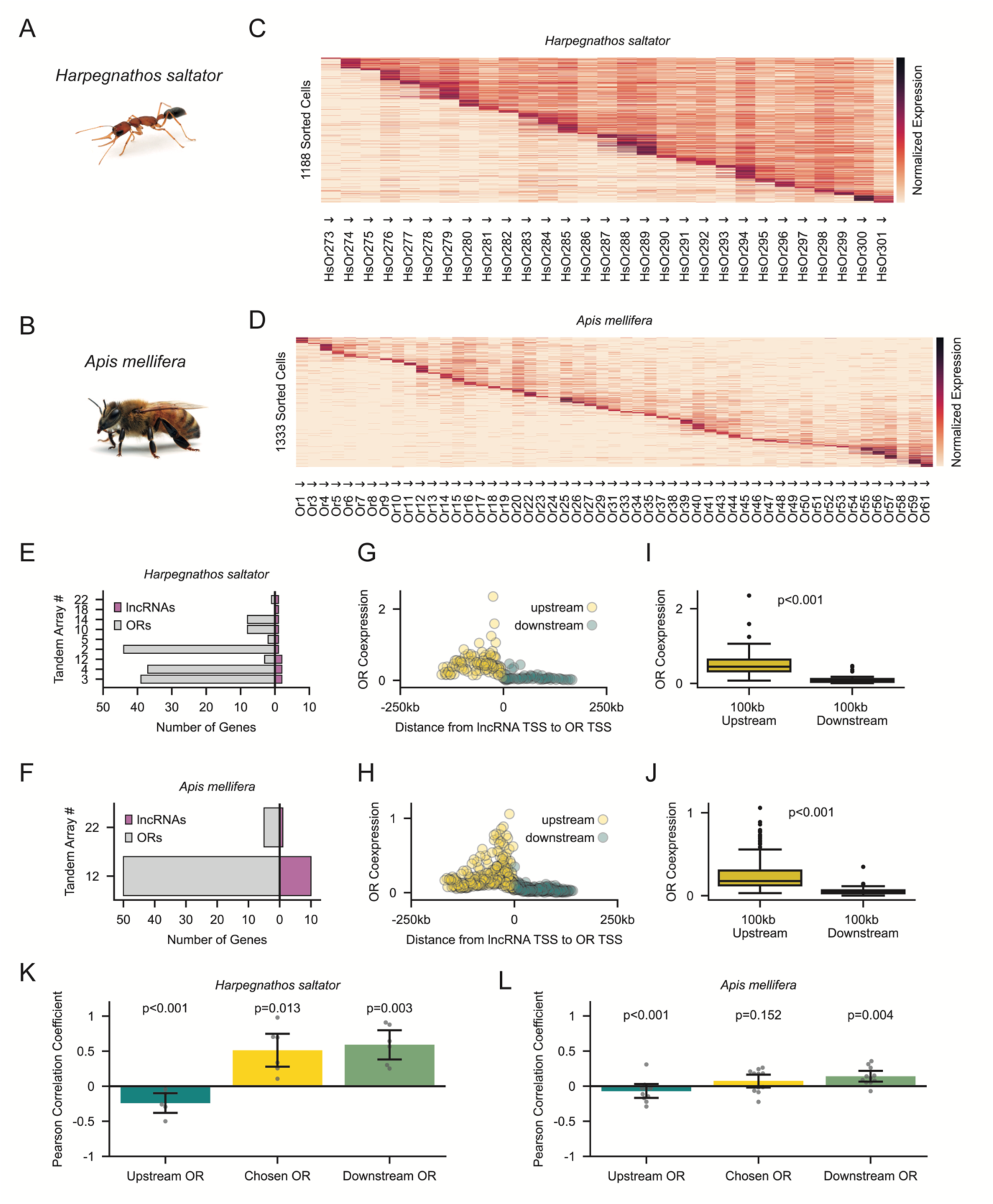
Bidirectional Promoter Activity in Other Ants and Bees, Related to Figure 5. (A-B) Photographs of *Harpegnathos saltator* (A) and *Apis mellifera* (B) workers (images by Alex Wild). (C-D) Representative tandem arrays from *H. saltator* (C; 30 ORs) and *A. mellifera* (D; 53 ORs). Heatmaps of log-normalized expression of all ORs in each tandem array across cells with a chosen OR in the corresponding tandem array. Cells (rows) are sorted by the genomic position of their chosen OR. Arrows indicate strand orientation. (E-F) Number of antisense-annotated lncRNAs (magenta) nested within each tandem array and the corresponding number of ORs per array (grey) in *H. saltator* (E) and *A. mellifera* (F). (G-H) Mean log-normalized coexpression of upstream (yellow) and downstream (cyan) ORs vs. the TSS-TSS distance from lncRNAs, using antennal snRNA-seq data from *H. saltator* (G) and *A. mellifera* (H). Each dot represents a cell in which the corresponding lncRNA is detected. (I-J) Boxplots of log-normalized OR coexpression within 100 kbp upstream (yellow) or downstream (cyan) of nested antisense lncRNAs, using antennal snRNA-seq data from *H. saltator* (I) and *A. mellifera* (J). Each boxplot shows the median and quartiles; the whiskers extend to 1.5 times the interquartile range. P-values from Wilcoxon rank-sum tests. (K-L) Pearson correlation coefficients for each unique lncRNA and either upstream ORs (cyan), chosen ORs (yellow), or downstream ORs (green), using antennal snRNA-seq data from *H. saltator* (K) and *A. mellifera* (L). Each lncRNA has a 3’ end within ≤5 kb of the chosen OR TSS. P-values from one-sample t-tests against zero. Error bars: 95% CI centered on the mean.

**Figure S7.**
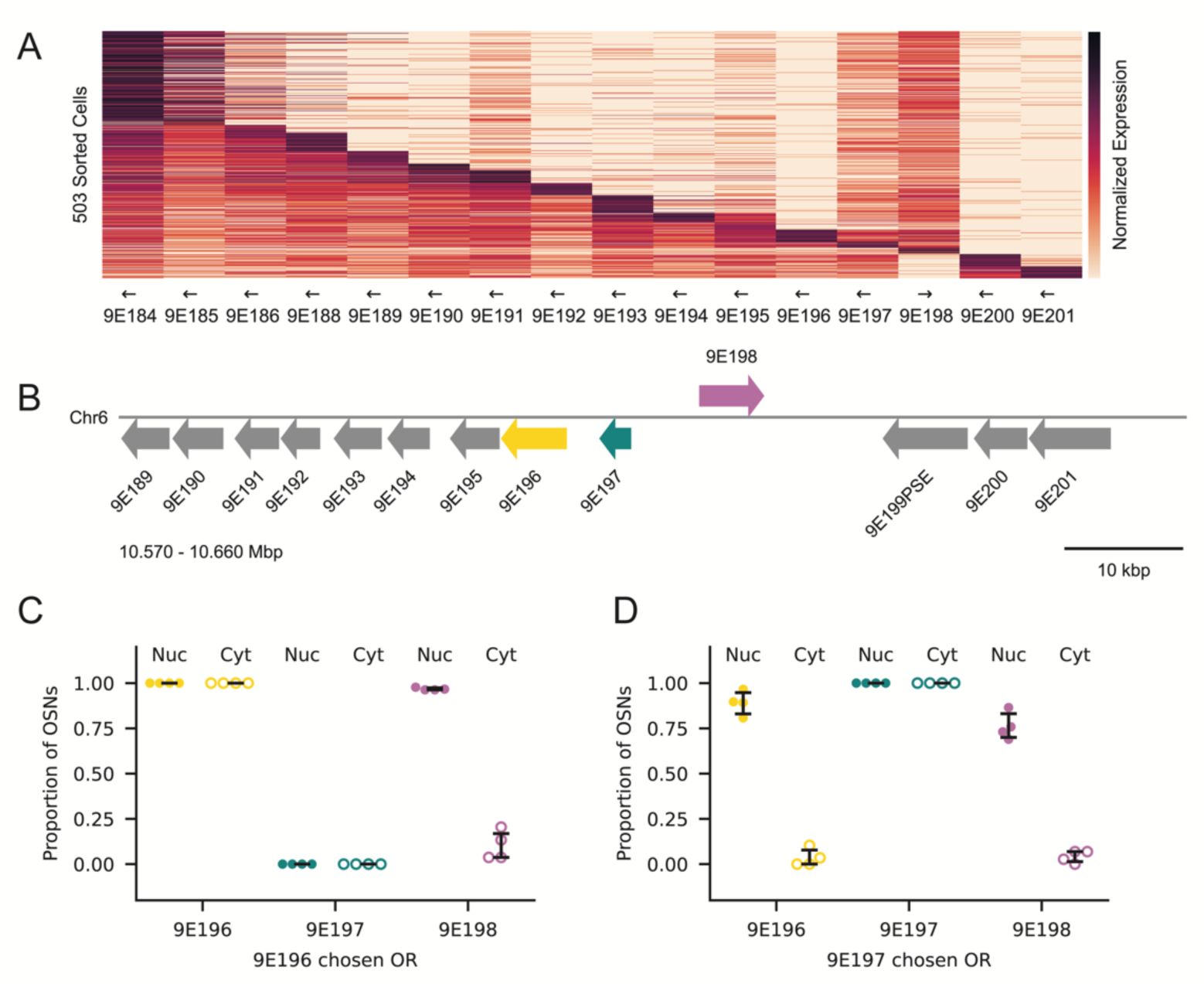
Additional Staining of Inverted OR Genes, Related to Figure 7. (A) Heatmap of log-normalized expression of all ORs in T35 across cells with a chosen OR in T35. Arrows indicate strand orientation. (B) Schematic of a subset of T35, highlighting ORs 9E196 (yellow), 9E197 (cyan), and 9E198 (magenta). (C–D) Proportion of OSNs with 9E196 (C) or 9E197 (D) as the chosen OR per antenna (n=4) exhibiting 9E196, 9E197 and 9E198 signal in the nucleus and cytoplasm. Error bars: 95% CI centered on the mean.

